# Precocious maturation and Purkinje cell dysfunction in the mouse *Chd8* haploinsufficient cerebellum

**DOI:** 10.64898/2026.09.26.754669

**Authors:** Nicolas Seban, Esteban Merino, Alexa D’Ambra, Se Jung Jung, Jade Lin Dungca, Stephanie A. Lozano, Ethan Fenton, Nickolas Chu, Kiya Jackson, Darlene Rahbarian, Aspen Kopley-Smith, Melissa Corea, Karol Cichewicz, Cory Ardekani, Ayanna Wade, Evangelos G. Antzoulatos, Cesar P. Canales, Diasynou Fioravante, Alex S. Nord

## Abstract

The cerebellum plays an emerging, but understudied, role in neurodevelopmental disorders (NDDs). Mutations in *CHD8* cause a monogenic NDD (CHD8-NDD) characterized by autism and other manifestations, and prior work in *Chd8* mutant mice reported cerebellar abnormalities, raising the possibility of cerebellar contribution to CHD8-NDD. To characterize the impact of *Chd8* haploinsufficiency on cerebellar development and function, we profiled the postnatal day 12 (P12) cerebellum of germline *Chd8^+/-^*mice using transcriptomic, morphological, and electrophysiological approaches. Gross cerebellar morphology was unaffected, but single-nucleus RNA sequencing revealed transcriptomic signatures of pathophysiology across cell types, including accelerated maturation of molecular layer interneuron, granule cell, and oligodendrocyte lineages. Purkinje cells (PCs) showed the strongest transcriptomic burden of any cell type and, consistent with this, exhibited altered morphology and electrophysiological properties. Bulk RNA sequencing confirmed that transcriptomic pathology persists into adulthood, indicating lasting impacts on both neuronal and glial gene expression. Together, these results capture cell-type-specific cerebellar perturbations associated with *Chd8* haploinsufficiency and support the cerebellum as a relevant locus of pathology in CHD8-NDD.

---

The cerebellum (CB) is traditionally viewed as a motor control center, but emerging evidence highlights its critical role in social and cognitive behavior^1–3^. Reflecting this expanded role, CB dysfunction has been linked to cognitive, affective, and social disorders in animal models and human studies^4–7^, and is increasingly recognized as a key contributor to neurodevelopmental disorders (NDDs), particularly autism spectrum disorder (ASD). Human imaging studies frequently identify CB dysmorphology in individuals with ASD^8^, and mouse models have directly connected monogenic NDDs to CB pathology. For instance, mutations in high-risk NDD genes such as *Tsc1* and *Shank2* disrupt CB development, impair CB circuit function and perturb key CB cell types, including Purkinje cells (PCs), the primary output neurons of the cerebellar cortex^9–11^. By identifying molecular, cellular, and circuit-level phenotypes in the CB, these studies expand our conceptual framework of NDD etiology and reshape our view of CB contributions to brain disorders.

Among monogenic NDD risk genes, the chromatin remodeler Chromodomain Helicase DNA-binding protein 8 (*CHD8*) stands out due to its high frequency of de novo loss-of-function mutations and high ASD penetrance^12,13^. Individuals with *CHD8*-associated NDD (CHD8-NDD) carry single allele loss-of-function mutations of the *CHD8* gene and present with a broad pathology spectrum in addition to ASD, including intellectual disability, macrocephaly, sleep and gastrointestinal dysfunction, and psychiatric comorbidities^14–20^, with some evidence for altered CB structure. Germline and conditional *Chd8* mouse models have recapitulated key NDD-relevant phenotypes^21–23^, yet how *Chd8* haploinsufficiency impacts different brain regions and specific cell types remains incompletely understood. Macrocephaly in *Chd8* mutant mice, for example, is widely attributed to expanded cerebral cortical volume and the large majority of *Chd8* mouse studies have focused on the forebrain^21,23,24^, but whether other brain regions critical for cognitive and social behavior are similarly impacted remains an open question. This question is made more pressing by our previous observation that the deep cerebellar nuclei (DCN), the sole excitatory output of the CB^1^, show volume changes in the *opposite* direction in adult *Chd8* haploinsufficient mice (i.e., decreased volume)^21^, raising the possibility that the CB is an underappreciated component of CHD8-NDD.

The impact of germline *Chd8* haploinsufficiency on the developing CB is understudied, even though *Chd8* is expressed in the developing and adult CB^25,26^. While homozygous embryonic loss is lethal, lineage-specific homozygous conditional deletion of *Chd8* in CB granule cell progenitors (GCPs) disrupts progenitor proliferation and differentiation, alters CB lamination, and causes CB hypoplasia and motor coordination deficits. By contrast, neither conditional deletion restricted to postmitotic PCs nor lineage-restricted heterozygous conditional deletion in GCPs produced overt phenotypes^27,28^. Although valuable from a fundamental biology perspective, isolating the effects of complete, homozygous *Chd8* loss within individual cell lineages does not capture the systemic, dosage-sensitive, and disease-relevant effects of *Chd8* germline heterozygous mutation, which is the mutational context that models human CHD8-NDD. Consequently, and despite clear evidence that *Chd8* is required for CB development, how germline *Chd8* haploinsufficiency shapes cell-type-specific transcriptomic landscapes, morphological maturation, and circuit function in the developing CB remains unknown.

To address this gap, we characterized the CB of germline *Chd8* heterozygous mice at postnatal day 12 (P12) using an integrated molecular, morphological, and electrophysiological approach (**Fig. 1A**). We combined broad histological profiling with single-nucleus RNA sequencing (snRNA-seq) to resolve anatomical and cell-type-specific pathology, evaluated PC morphology and electrophysiology to determine if transcriptional perturbations correspond to functional deficits, and compared bulk RNA-seq profiles between P12 and P60 to establish whether molecular pathology is transient or persists into adulthood. Together, our results define cellular, regional, and temporal consequences of NDD-relevant *Chd8* haploinsufficiency in the developing CB.

**Figure 1:**
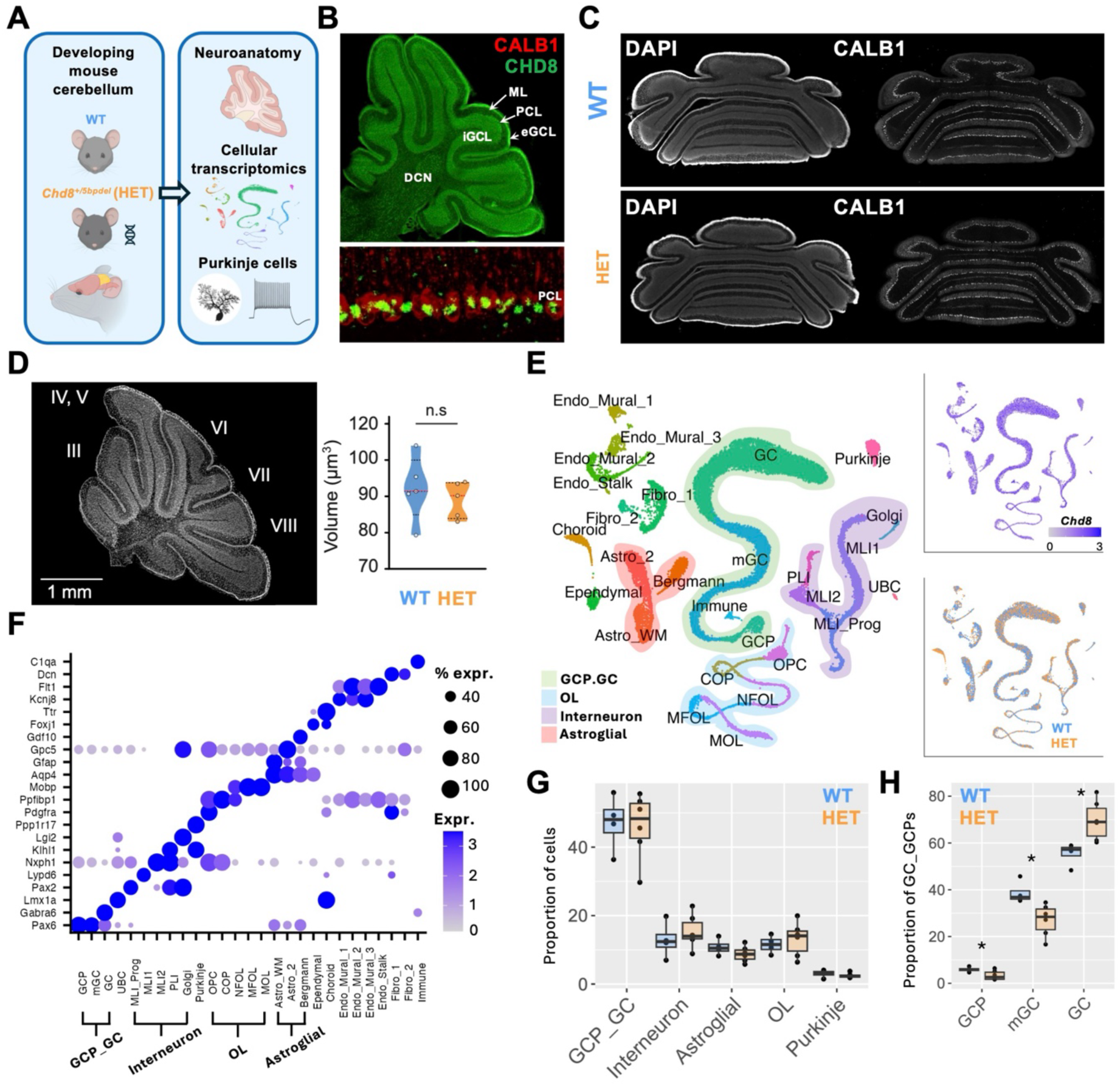
Developing *Chd8^+/5bpdel^*mice do not display gross anatomical or major cellular differences in the cerebellum. **A**) Schematic overview of this study. **B**) Sagittal CB section immunostained for CHD8. CB anatomical divisions are highlighted to show ubiquitous expression. Molecular layer (ML), Purkinje cell layer (PCL), internal and external granule cell layer (iGCL, eGCL), and deep cerebellar nuclei (DCN). A zoomed in image of the PCL is co-stained for CALB1, a PC marker. **C**) Example WT and *Chd8^+/5bpdel^* coronal sections stained with DAPI and CALB1 to examine broad CB anatomy, demonstrating typical formation of the CB cortex in *Chd8^+/5bpdel^* mice. **D**) Representative sagittal CB section and quantification of total CB volume (summed across lobules), showing no significant differences between genotypes. **E**) UMAP of 21,463 CB nuclei from P12 WT and *Chd8^+/5bpdel^* littermates with cell types annotated. Shading highlights cell type supertypes composed of similar or developing lineages (Astroglial, Interneurons, OL, GCP_GC). Inlayed UMAPs show ubiquitous expression of *Chd8* in CB cells and confirmation that WT (n = 5) and *Chd8^+/5bpdel^* (n = 6) cells intermix within cell type clusters. **F**) Cell type identification dot plot representing average expression of marker genes. Supertype annotation of cell types is also shown. **G**) Cell type proportion comparison between *Chd8^+/5bpdel^* and WT nuclei show no significant differences across cell supertypes using Student’s t-test. **H**) Proportion of developing granule cells show significant differences via Student’s t-test for granule cell progenitors (GCP) (p = 0.034), migrating granule cells (mGC) (p = 0.013), and differentiated granule cells (GC) (p= 0.010).

## Results

### Gross cerebellar morphology and cortical lamination are spared in *Chd8* haploinsufficient mice

To examine the impact of *Chd8* haploinsufficiency on the developing CB, we utilized our previously published germline *Chd8* heterozygous loss-of-function mouse model (*Chd8^+/5bpdel^*). *Chd8^+/5bpdel^*mice recapitulate macrocephaly associated with *CHD8*-NDD and display subtle learning and memory deficits^21^. Building on our previous finding of decreased DCN volume in adult *Chd8^+/5bpdel^*mice, here we examined the CB at P12, a landmark stage in CB development marked by eye opening and the onset of sensorimotor integration^27,28^. This time point captures ongoing postnatal neurogenesis of granule cells (GC) and molecular layer interneurons (MLI), as well as the active dendritic maturation and synaptic refinement of PCs and other resident cell types^29^.

To confirm CHD8 protein expression in the developing CB, we performed immunohistochemistry (IHC) in P12 wildtype mice. Sagittal CB sections showed CHD8 expression across all major CB subregions (**Fig. 1B**): molecular layer (ML), Purkinje cell layer (PCL), external and internal granule cell layer (eGCL, iGCL), white matter (WM), and deep cerebellar nuclei (DCN). To test whether *Chd8* haploinsufficiency alters CB morphology, we examined sagittal and coronal sections for gross alterations in lobule structure and laminar patterning. IHC stains for Calbindin 1 (CALB1), co-stained with DAPI nuclear marker, allowed visualization of overall CB lamination and structure. Compared with WT, *Chd8^+/5bpd^*^el^ sections showed no obvious malformations or disruption to the ML, PCL, eGCL, iGCL, or DCN (**Fig. 1C**, **Supplemental Fig. 1A,B**), nor any evident difference in PC monolayer formation (P = 0.44, n = 6 WT, n = 6 *Chd8^+/5bpdel^*; **Supplemental Fig. 1C,** see **Supplemental Table 1** for details on statistical methods). Serial sagittal sections were also used to quantify total CB cortex volume (WT and *Chd8^+/5bpdel^* = 18 slices from 5 mice per group, both sexes) and no significant difference was detected between genotypes (t_8_ = 0.669, p = 0.522) (**Fig. 1D**). Together, these results indicated that CHD8 is broadly expressed across CB, and that germline *Chd8* haploinsufficiency does not produce gross alterations to CB morphology and/or lamination at P12.

### Cell-type-specific perturbation of developmental trajectories and gene expression profiles in *Chd8^+/5bpdel^*cerebellum

To examine CB cellular and molecular changes in *Chd8^+/5bpdel^*mice with cell-type-specific resolution, we performed snRNA-seq on tissue from 5 WT (both sexes) and 6 *Chd8^+/5bpdel^* (both sexes) mice at P12 (see Methods for full experimental and computational details). We generated data for 21,463 nuclei that passed quality-control filtering, with an average of 7,308 unique RNA counts and 2,555 gene features per cell (**Supplemental Fig. 2A-C)**. Following dimensionality reduction and clustering, we identified 62 clusters mapping to CB cell types, distributed across both discrete and continuous population distributions (**Fig. 1E**, **Supplemental Fig. 2D,E**). Using cell-type-specific gene markers^30^, we identified all expected CB cell types and developmental lineage trajectories in P12 CB cortex (**Fig. 1F**). Our dataset included 6 GABAergic cell types: PCs; Golgi interneurons (GoC); Purkinje layer interneurons (PLI); two MLI subtypes: MLI-1, which synapse onto PCs, and MLI-2, which synapse onto MLI-1; and MLI progenitors, which remain in the WM until differentiated into MLI-1 or MLI-2 subtypes^31^. We also identified glutamatergic interneurons including unipolar brush cells (UBC), which integrate signals within the GCL; and GCs, which form parallel fibers and provide excitatory input to PCs alongside climbing and mossy fibers originating outside the CB^11,32^. GCs constitute 99% of the neurons in the mature mouse CB cortex^33^, but are still expanding at P12, making up 47% of cells in our dataset and spanning three major developmental states: proliferative granule cell progenitors (GCP), migrating granule cells (mGC) originating from eGCL, and terminally differentiated GCs settled in iGCL.

We identified three major glial classes: white matter astrocytes (Astro_WM), astrocytes located primarily in grey matter regions (Astro_2), and CB-specific Bergmann glia residing in the PCL. A large cluster of differentiating oligodendrocytes (OL) was resolved by developmental state^34^, comprising proliferating oligodendrocyte precursor cells (OPCs), terminally committed oligodendrocyte precursors (COP), newly formed oligodendrocytes (NFOL) extending processes toward neuronal axons, myelinating oligodendrocytes (MFOL) actively generating myelin sheaths, and mature oligodendrocytes (MOL) with established myelin sheaths. We additionally identified a cluster of neuroectoderm-derived ependymal cells, a glial type supporting vasculature; four endothelial cell types, including three mural cell clusters (Endo_Mural_1/2/3) and stalk cells (Endo_Stalk) that line and regulate CB blood vessels^35^; two fibroblast populations (Fibro_1/2); immune cells comprising macrophages and microglia; and choroid plexus cells. Finally, we identified a cluster of presumed projection neuron (PN) subtypes, including DCN PNs as well as non-CB midbrain and hindbrain PN classes captured incidentally in the dissection (**Supplemental Fig. 4A,B**).

Major cell types (PCs, GCs, GABAergic interneurons, astroglia, and oligodendrocytes) were well represented across sex and genotype (**Fig. 1E**, **Supplemental Fig. 2F,G**). Likely reflecting proximity to vasculature and ventricles bordering the dissection site, endothelial (Mural, Stalk), immune, and endothelial-like (Choroid, Fibroblast, Ependymal) clusters showed high variance across replicates regardless of genotype and were excluded from downstream analyses (**Supplemental Fig. 2H**). Cell types were grouped by superclass into Cell Type Level 1 (CT1: OL, Interneuron, GCP_GC, Astroglial, PC) to assess effects within broader populations.

We then tested for altered CB cell proportions and cell-type-specific molecular pathology associated with germline *Chd8* haploinsufficiency. We first tested for differences in the abundance of cell types between WT and *Chd8^+/5bpdel^*P12 CB by comparing library proportion for each cell population across biological replicates with Student t-test (see **Supplemental Table 1** for statistics details for all transcriptomics analyses). Consistent with gross anatomy, we found no significant differences in the number of CT1 CB cortex cell classes (GCP_GC, Interneurons, Astroglial, OL, PC) at the level of nominal P values or those corrected for multiple testing (**Fig. 1G**). At the individual cell type level (**Supplemental Fig. 3A**), we observed decreases in developing populations, including GCP (p = 0.028) and OPC (p = 0.055) clusters, though these were not significant with multiple-testing corrections. OPCs also showed a significant increase in the proportion of dividing (in G2M phase) cells in *Chd8^+/5bpdel^* samples compared to WT (p = 0.048; **Supplemental Fig. 3D**). We next assessed relative abundance of state-associated cell classes within lineages where a developmental trajectory was captured (GCs, OLs, and MLIs) (**Supplemental Fig. 3C**). In all three lineages, we found significant increases in mature cell states and decreases in immature states. For example, within the GC lineage (**Fig. 1H**), we found decreased progenitor (p = 0.034) and migrating (p = 0.013) populations alongside increased terminally differentiated clusters (p = 0.010) expressing the mature marker *Gabra6*. Finally, although we could not resolve specific classes, we observed an overall decrease in PN proportion in *Chd8^+/5bpdel^* mice (**Supplementary Fig. 4C**). Together, these cell proportion analyses indicate no differences in major CB cell classes, but altered developmental composition within multiple cell lineages and a decrease in general projection neurons in *Chd8^+/5bpdel^* mice.

We next asked whether global transcriptional signatures within any cell types were sufficient to distinguish *Chd8^+/5bpdel^*from WT CB, applying scDist, a linear mixed-effects framework that estimates condition-associated shifts in high-dimensional gene-expression space while accounting for biological variability^36^. PCs were the only population exhibiting a significant transcriptomic shift between *Chd8^+/5bpdel^* and WT mice after multiple-testing correction (FDR = 0.084; **Supplemental Fig. 3B**). This result identifies PCs as strongly impacted by *Chd8* haploinsufficiency at the transcriptomic level, an intriguing finding considering the central role PCs play in integrating CB cortical signaling.

In summary, snRNA-seq of the P12 CB resolved the major neuronal and glial populations and their developmental trajectories in WT and *Chd8^+/5bpdel^*mice. Multiple developing lineages, including GCs, oligodendrocytes, and MLIs, showed a shift from progenitor or immature states toward more mature states. Complementary cell prioritization analysis identified PCs as showing the strongest genotype-associated transcriptomic perturbation. Together, these findings indicate that *Chd8* haploinsufficiency alters CB developmental progression and drives prominent cell-type-specific molecular impacts in PCs.

### Accelerated pseudotime signatures in CB lineages of P12 *Chd8^+/5bpdel^* mice

Observed changes in cell type proportions in GC, interneuron, and oligodendrocyte lineages appeared to be driven by altered developmental progression. To test this directly, we examined pseudotime as implemented in Monocle3^37^, analyzing the relative rate of development for cells within each lineage. While developmental effects may be present in other cell types, only these three lineages could be assessed using a pseudotime framework, as other populations did not show sufficient spread across developmental states at P12. Pseudotime analysis was performed on each developing cell type, anchored on the progenitor state. Pseudotime scores were calculated based on nearest neighbors across all WT and *Chd8^+/5bpdel^*cells combined, with correct trajectory ordering confirmed by the expected spread of scores across immature and mature cell states. Differences between WT and *Chd8^+/5bpdel^* cells and samples were analyzed both across all cells in each lineage, and within individual developmental states, with genotype-level differences determined by Student t-test comparing median pseudotime score across biological replicates. Monocle3 was used to identify genes varying along pseudotime, applying Moran’s test statistic to detect nonrandom spatial patterns of gene expression across the inferred trajectory and assign genes into immature or mature gene sets (**Supplemental Fig. 5A, Supplemental Table 2**). DESeq2 pseudobulk Wald Test was used to assess differential expression of these developmental gene sets, with significance determined by FDR-corrected Wilcoxon rank sum test. Gene set enrichment analysis (GSEA) was implemented via the ClusterProfiler^38^, ranking developmental genes by Moran’s test statistic. When possible, developmental transcriptional state phenotypes were validated against external longitudinal single-cell datasets in the CB. This framework revealed significant developmental perturbation in each of the three lineages (**Fig. 2**).

**Figure 2:**
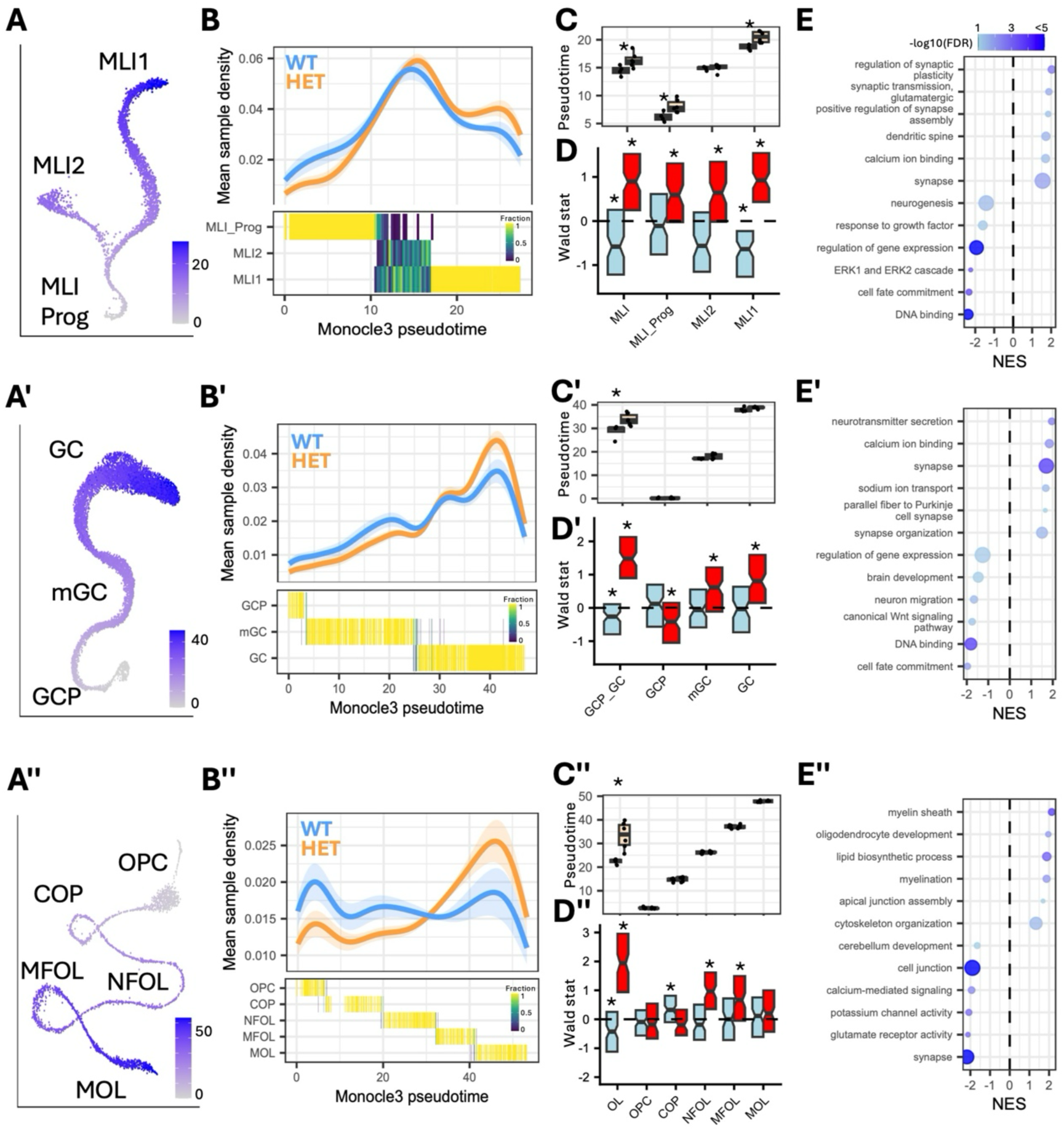
Newly born cell types in the P12 cerebellum show accelerated maturation in *Chd8^+/5bpdel^*mice. **A**) UMAP visualization of developing lineages colored by their Monocle3 pseudotime score: A. MLI, A’ Granule Cells (GC), A’’. Oligodendrocytes (OL). **B**) Upper, density plot of pseudotime scores across WT and *Chd8^+/5bpdel^* (HET) samples, standard error across biological replicates is shown with shading. Lower, heatmap of pseudotime score across developing lineages. B. MLI cell types showing pseudotime score captures progression of MLI progenitors (MLI_Prog) to MLI-1 and MLI-2. B’. GC cell types showing pseudotime score captures progression of granule cell progenitors (GCP) to migrating granule cells (mGC) to differentiated granule cells. B’’. OL cell types showing pseudotime score captures progression of oligodendrocyte progenitors (OPCs) to mature OLs (MOL). **C**) Boxplot showing median pseudotime score across biological replicates across entire developmental lineage and individual subtypes, significance tested by Student’s t-test. C. Significant differences were found across entire MLI lineage (p = 0.026), MLI_Prog (p = 0.019), and MLI1 (p = 0.007). C’. Significant differences were found across the entire GC lineage (p = 0.03), but not in subtypes. C’’. Significant differences were found across entire OL lineage (p = 0.004) but not in subtypes. **D**) Boxplot showing Wald-stat distributions derived from DESeq2 for genes correlated with high (red) and low (blue) Pseudotime scores in the developing trajectories. * indicates FDR< 0.05 differences between genotypes. D. MLI D’. GC. D’’. OL. **E**) Significant (FDR < 0.05) GO terms associated with pseudotime genes derived from GSEA analysis. E. MLI. E’. GC. E’’ OL.

MLIs are the last GABAergic cell type born in the CB, with progenitors continuing to proliferate in the WM through the second postnatal week^29,39^. Maturing MLIs then migrate into the ML and terminally differentiate into either MLI-1 or MLI-2 classes. MLI-1 cells are born earlier and directly synapse onto PCs, whereas MLI-2 are born later and form synapses onto MLI-1^31^. Pseudotime analysis captured this trajectory in our dataset (**Fig. 2A**), with scores across MLI developmental trajectory shifting toward higher pseudotime values in the *Chd8^+/5bpdel^* mice relative to WT littermates (**Fig. 2B**). Significant differences were found for the full lineage (p = 0.026), as well as for MLI_Prog (p = 0.019), and MLI-1 (p = 0.007). Genes associated with mature pseudotime state were increased across all MLI subtypes and the trajectory as a whole, with the reverse pattern for immature state-associated genes (**Fig. 2D**). Mature state-associated genes were enriched for terms associated with mature neuron function such as *synapse*, *calcium ion binding*, and *dendritic spine*. Notably, mature MLI genes were enriched for proteins functioning in postsynaptic and glutamatergic synapses, suggesting that maturation of postsynaptic, rather than presynaptic, machinery is preferentially accelerated. Genes associated with immature state were enriched for earlier developmental terms such as *DNA binding*, *neurogenesis*, and *cell fate commitment* (**Fig. 2E**). The pseudotime model derived from our MLI data was replicated in an external single-cell dataset^30^, supporting the robustness of our developmental model (**Supplemental Fig. 5B**). While MLI-2 neurons did not show a significant difference in the pseudotime score, they did show increased expression (FDR < 0.001) of mature neuronal gene sets (**Fig. 2D**). Together, these results indicate accelerated development of MLIs in *Chd8^+/5bpdel^* mice and increased expression of genes associated with mature MLI function.

GCs are the other neuronal cell type undergoing active neurogenesis in the P12 CB: GCPs proliferate in the eGCL, and post-mitotic mGC then migrate into the iGCL to form synaptic connections with PCs^29^. At P12, we capture this entire trajectory, with pseudotime scores tracking the transition from GCPs to mGC to differentiated GCs (**Fig. 2A**’). Overall, GC lineage pseudotime scores were higher in *Chd8^+/5bpdel^* mice (**Fig. 2B’**), with a significant difference between the median scores across the entire GC lineage (p = 0.03) and similar trends within individual GC cell states (**Fig. 2C’**). We observed strong differential expression of pseudotime marker gene sets across the lineage as a whole, along with upregulation of mature genes specifically within the migrating and differentiated states (**Fig. 2D’**). As with MLIs, genes associated with mature state were enriched for neuronal terms, while immature genes were enriched for earlier developmental terms (**Fig. 2E’**). Mature GC genes also showed strong enrichment for *neurotransmitter secretion*, suggesting increased excitatory drive onto afferent targets such as PCs. We validated the GC lineage pseudotime model in an external dataset covering longitudinal GC development^40^ (**Supplemental Fig. 5C**). These results indicate a shift toward a more differentiated state for GCs *in Chd8^+/5bpdel^* mice, with fewer GCP and mGC cells and increased mature GCs and parallel fiber synapses onto PCs. Unlike MLI-1, where we observed strong pseudotime differences in the mature population, the differentiated GC population itself does not show a large pseudotime shift, though it does show increased expression of mature state genes.

The final lineage captured by pseudotime was OLs. Unlike CB neurons, OPCs continue to proliferate and differentiate into mature OLs well into adulthood^41^. Our P12 data effectively recapitulated the full span of OL development, from OPCs, which transition to COPs, to NFOLs attaching to axons and initiating myelination programs, to MFOLs actively myelinating neuronal axons, and finally MOLs with fully formed myelin sheaths (**Fig. 2A’’**). As with neuronal populations, we observed a shift towards more mature states in *Chd8^+/5bpdel^* mice (**Fig. 2B’’**). OLs showed a significant pseudotime difference only when assessed across the entire developmental trajectory (p = 0.004) (**Fig. 2C’’**). Similarly, the strongest differential expression of pseudotime genes was observed when considering all OLs together, indicating a broad shift in developmental state (**Fig. 2D’’**). NFOL and MFOLs showed the strongest upregulation of mature OL genes, suggesting this transition may be affected the most. Genes associated with mature pseudotime state were enriched for terms reflecting OL identity and function (e.g. *myelin sheath* and *lipid biosynthetic process*, **Fig. 2E’’**). Of note, immature OL genes showed strong enrichment for neuro-associated terms such as synaptic signaling and transporter activity (**Fig. 2E’’**), likely reflecting ion channel and cell-sensing genes expressed by OPCs and COPs as they differentiate and locate their neuronal partners^42^. These results highlight a shift in OL state marked by upregulation of the myelination machinery during OL maturation. Together, the pseudotime analyses indicate a consistent shift toward more advanced transcriptional maturation across the MLI, GC, and OL lineages in the P12 *Chd8^+/5bpdel^* CB.

### Cell-type-specific and shared differential gene expression signatures in *Chd8^+/5bpdel^* cerebellum

To characterize cell-type-specific transcriptional phenotypes, we performed differential expression (DE) analysis using a cluster-level pseudobulk framework, in which reads from each relevant cell type were aggregated at sample level and DE testing was performed with DESeq2, including covariates controlling for technical variation and sex. DE genes (DEGs) were defined as those with p-value < 0.05 and |log2 fold change| > 0.1, with full DE results reported in **Supplemental Table 3**. We first compared DE burden (the number of DEGs divided by the number of genes expressed in that cell type, only retaining protein coding genes) and enrichment for SFARI ASD-associated genes among DEGs (**Fig. 3A**). PCs had the strongest DE burden, with 4.2% of expressed genes (306 DEGs) differentially expressed. MFOLs were next, though with a sharp relative drop-off, at 3.3% of MFOL genes (172 genes) DE. PCs were also the only cell type showing significant enrichment for ASD-associated genes (FDR = 0.01). DEGs in each CB cell type were largely distinct, demonstrating the cell-type-specificity of these transcriptomic perturbations (**Fig. 3B**, **Supplemental Fig. 6A,B**). However, we did identify a set of genes with evidence for shared DE across cell types (**Supplemental Fig. 6C**).

**Figure 3:**
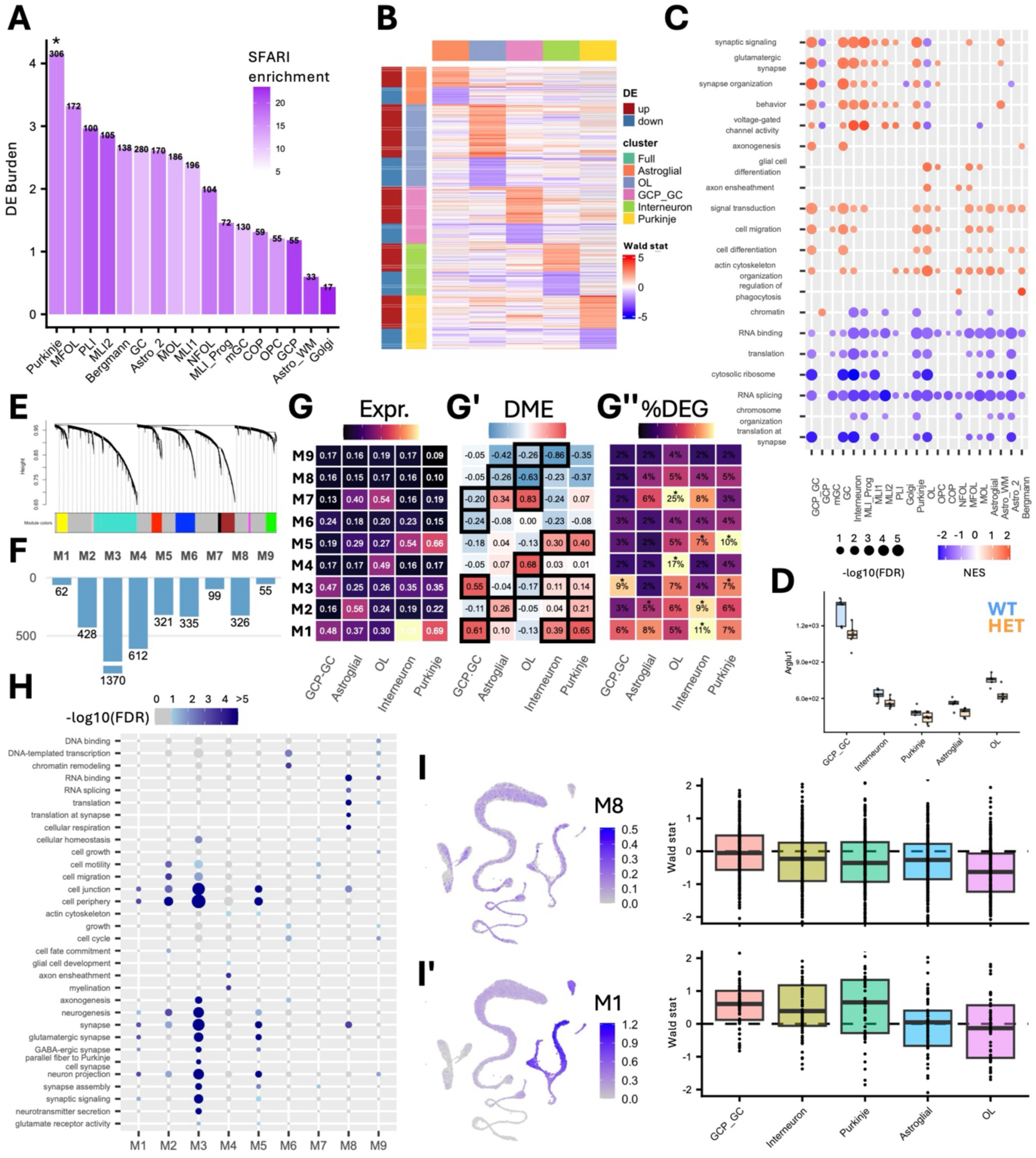
*Chd8^+/5bpdel^*transcriptional pathology highlights conserved perturbation to basal cellular processes across cell types and increased susceptibility of PCs. **A**) Bar plot showing differential expression (DE) burden across CB cell types (Y-axis value) and relative enrichment and count for ASD-associated genes (color intensity, numbers). DE burden is calculated as the percentage of expressed genes with significant (p < 0.05) DE from pseudobulk DESeq2 analysis. *FDR < 0.05 for SFARI gene enrichment. **B**) Heatmap showing the cell-type-specific DE across major CB cell type classes. Genes included have significant (p < 0.05) DE in at least one cell type and color indicates the DESeq2 Wald stat. **C)** Curated GSEA results from cell type-specific DE results. Only nominally significant (p < 0.05) results are shown, dot size indicates corrected p-value (FDR) and color the normalized enrichment score (NES). **D)** Boxplot showing normalized expression for *Arglu1*, which was downregulated across cell types. **E**) Dendrogram from hdWGCNA run on the major cell classes. **F**) Bar plot showing the size of the 9 modules identified in hdWGCNA. **G**) Heatmap showing normalized expression of modules in the major cell types, as calculated from Seurat’s ModuleScore function. **G’**) Heatmap showing median Wald stat for each module from DESeq2 DE analysis on major cell types. Black boxed cells have a significant change in logFC compared to the grey module (Wilcoxon rank sum test, FDR < 0.01). **G’’**) Heatmap testing for overrepresentation of DESeq2 DEGs (p <0.05) by module for each major cell type. Percent indicates percent of module genes that are DE Starred cells have a significant (FDR < 0.05) enrichment for DEGs via a permutation test. **H)** Curated GO enrichment for hdWGCNA modules. Dot size indicates relative number of genes belonging to that term and color the corrected p-value (FDR). **I)** Examples of differential expressed module (DEM), M8(I) and M1(I’). Featureplot of the relative expression of the module in the UMAP space, as calculated from Seurat’s AddModuleScore. Boxplot showing differential expression of the module across major cell types, relative expression values derived from DESeq2 Wald stat.

We next performed GSEA functional enrichment based on DE rank within each cell type using ClusterProfiler^38^ (**Fig. 3C, Supplemental Table 4**). All cell types showed some level of significant (p < 0.05) functional enrichment associated with their differential transcriptional signatures, with several annotation classes across cell types. Across neuronal populations, there was a shared upregulation of terms associated with mature neuronal function, which was particularly strong in PCs. Shared across both neuronal and glial cell types was downregulation of basal cellular processes, including transcription, RNA binding, RNA processing and translation. A representative gene from this functional group, *Arglu1*, showed a conserved downregulation in the *Chd8^+/5bpdel^*samples across major cell types (**Fig. 3D**). *Arglu1* is a dual-function nuclear regulator that acts as a transcriptional coactivator and modulator of pre-mRNA alternative splicing^43^. OLs had the most distinctive GSEA signature, showing downregulation of neuronal function-associated terms, reflecting the higher ion channel activity present in progenitor and immature states, alongside upregulation of OL-specific genes associated with *glial cell differentiation* and *axon ensheathment*. Overrepresentation analysis using significant DEGs (p < 0.05) from each cell type using ClusterProfiler produced similar functional enrichment (**Supplemental Fig. 7A,B, Supplemental Table 5**), with a large number of terms significantly enriched in OLs, GCs, and PCs. Together, these analyses identify PCs as the most transcriptionally affected cell type in the P12 *Chd8^+/5bpdel^* CB and reveal cell-type-specific perturbations and shared increases in neuronal maturation programs alongside decreases in RNA-processing and translational pathways.

As an alternative to single-gene DE analysis, we performed hdWGCNA^44^ to identify co-expression modules and test for differential expression at the module or network level, focusing on populations with sufficient cell counts (interneurons, astroglial, GCP_GC, OL) (**Supplemental Fig. 8, Supplemental Table 6**). hdWGCNA identified nine main modules along with a non-correlated “grey” module (**Fig. 3E**), with varying gene membership size (**Fig. 3F**). To determine whether modules represented networks marking specific cell identity, we used Seurat’s ModuleScore function^45^ to compare cell-type-specific expression across populations (**Fig. 3G**). Modules M6, M8, and M9 showed non-specific expression across cell types, whereas the remaining modules were composed of genes largely specific to particular classes: M7 was primarily expressed in glial cells, M5 in GABAergic cells, M4 in OL, M2 in astroglial cells, and M3 and M1 in neuronal cells. To test for differential expression of these modules, we used the DESeq2 results to assess shifts in the distribution of Wald test scores among module gene sets (**Fig. 3G’**) and overrepresentation of DEGs within modules (**Fig. 3G’’**). To assign functional meaning to modules, we performed GO analysis using clusterProfiler (**Fig. 3H**).

M8 and M9 (**Fig. 3I**), which showed uniform expression across cell types, were both conserved in downregulation and associated with basal cellular processes such as *DNA-templated transcription, chromatin remodeling, RNA binding,* and *translation*. The glia-enriched M7 was strongly upregulated in OLs, and enriched for terms including *cell motility* and *cell migration*. M6 was primarily downregulated in the GC and interneuron lineages and enriched for basal processes as well as terms associated with cell development. M5 was upregulated in interneurons and PCs and associated with GABAergic and synaptic neuronal terms. M4 was OL-specific, strongly upregulated in OLs, and enriched for *axon ensheathment* and *myelination*. M3, the largest gene set, showed enrichment in neurons and upregulation within the GC lineage, with enrichment for terms associated with glutamatergic synapses and, notably, *parallel fiber to PC synapse*. M2 was enriched for a broader set of terms, including *cell periphery, nervous system development, cell migration*, and *signaling*, and was upregulated in astroglial cells and PCs. M1 (**Fig. 3I’**) showed strong enrichment and upregulation across all neuronal cell types and was associated with *synapse*, *neuron projection*, and *neurogenesis* terms.

Together, hdWGCNA revealed coordinated downregulation of broadly expressed transcriptional and translational modules alongside cell-type-specific activation of functional programs, including GABAergic synaptic genes in inhibitory neurons, glutamatergic and parallel fiber-to-PC synaptic genes in GCs, and myelination-associated genes in OLs. In summary, *Chd8* haploinsufficiency caused cell-type-specific transcriptomic dysregulation across CB cell types, with the largest and most prominent effect in PCs. While DEGs were largely cell-type-specific, we identified overlapping functional enrichment for synapse-associated gene ontology (GO) terms across neuronal cell types, alongside conserved downregulation of genes associated with basal cellular processes across all cell types, including transcription and translation.

### Transcriptional pathology associated with synaptic function and morphology in *Chd8^+/5bpdel^* Purkinje cells

Given that PCs are responsible for relaying CB cortical signals to downstream targets and exhibited the strongest DE burden in our analyses, we focused on transcriptomic effects of *Chd8* haploinsufficiency in these cells. Despite the relatively low number of PCs (556 cells) in our data set, this population featured the most DEGs (p < 0.05) of any cell type, with 131 downregulated and 202 upregulated genes (**Fig. 4A**). A small set of 20 genes passed the stringent FDR < 0.05 cutoff, including *Etl4*, *Gpc6*, and *Garnl3*. Rank-based GSEA analysis identified significant (FDR < 0.05) enrichment for upregulation of gene sets including *GABAergic synapse*, *glutamatergic synapse*, *neuron projection*, *action potential*, and *potassium ion transport*. Downregulated GSEA terms were associated with basal cellular processes discussed above, including *gene expression*, *chromosome organization*, *translation*, and *RNA splicing* (**Fig. 4B**). GO analysis was largely concordant with these GSEA results (**Supplemental Fig. 7A,B**). We visualized the annotation of DEGs to representative enriched neuronal GO terms, allowing us to examine gene overlap across synaptic functional annotations (**Fig. 4C**). Some DEGs were annotated to multiple terms, yet a large proportion was unique to specific neural GO processes, suggesting impacts on distinct neuronal functions. While the majority of DEGs within these gene sets was upregulated, some were downregulated, indicating a more complex pattern of perturbation.

**Figure 4:**
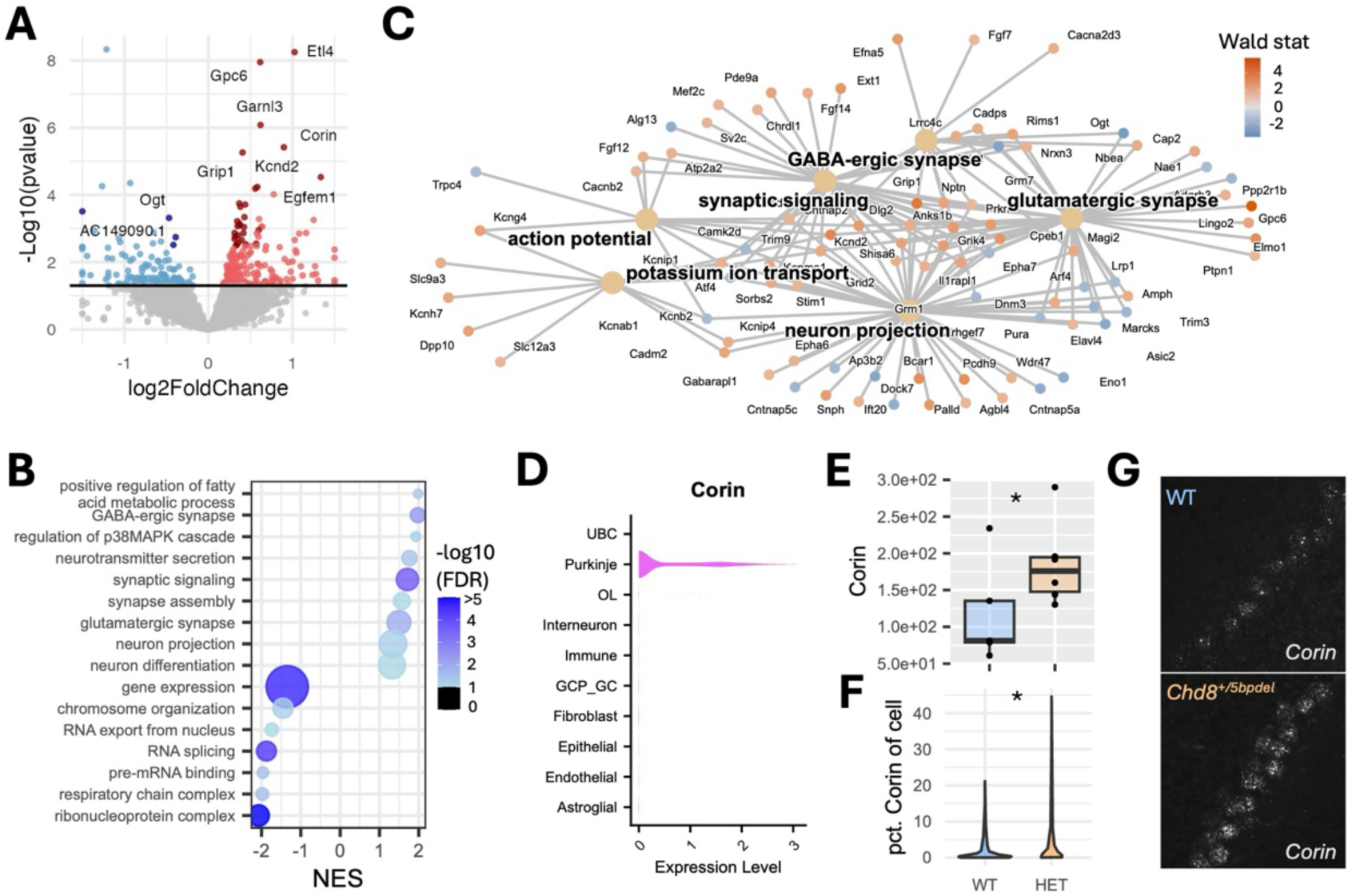
Purkinje cells feature transcriptomic perturbations to neuronal function and morphological processes. **A**) Volcano plot showing the DE results from pseudobulk DESeq2 in PCs. Color indicates significance of DE effect, light red (up, p < 0.05), dark red (up, FDR < 0.1), light blue (down, p < 0.05), dark blue (down, FDR < 0.1). **B**) Dot plot showing highlighted GSEA enrichment from PC DE results. Size indicates gene set size and color significance. **C)** Gene–concept network (Cnet) plot showing the mapping of DEGs (P < 0.05) to significant GO terms enriched in Purkinje cell DE (FDR < 0.05). Color reflects the DESeq2 Wald stat. **D**) Violin plot showing PC specific expression of *Corin*. **E)** Boxplot showing differential expression of *Corin* in PCs (FDR < 0.001). **F**) Violin plot showing quantification of per-cell Corin signal from RNA FISH in WT vs *Chd8^+/5bpdel^* PCs, pooled across CB lobules in coronal sections. Significance determined by Wilcoxon test (p < 0.001). **G**) Representative *Corin* RNA FISH images from PCs.

To validate the cell-type-specific DE identified in PCs, we selected *Corin* as a candidate gene, given its strong significance (FDR < 0.001) (**Fig. 4E**) and PC-specific expression (**Fig. 4D**). *Corin* is a transmembrane serine protease previously shown to be upregulated in certain classes of adult PCs^46,47^, although its role in developing PCs is unknown. We used RNA fluorescence in-situ hybridization (FISH) via the RNAScope platform to label *Corin* mRNA in P12 serial CB coronal sections of WT (n= 10) and *Chd8^+/5bpdel^* (n= 10) mice (both sexes), with PCs co-labeled using an antibody targeting CALB1. Sampling across multiple lobules across the CB, we surveyed 1,164 WT and 1,192 *Chd8^+/5bpdel^* PCs. *Corin* was not expressed in all PCs, suggesting it may mark a specific cellular state or subtype. Comparing WT and *Chd8^+/5bpdel^*PCs, we found a significant increase in both the proportion of PCs expressing *Corin* (p = 0.041 generalized linear model) and the level of *Corin* expression (p < 0.001, Wilcoxon rank-sum test) (**Fig. 4F,G**). Together, these transcriptional findings identify PCs as a major axis of transcriptional pathology in the *Chd8^+/5bpdel^* CB.

### Altered cellular morphology and function in *Chd8^+/5bpdel^* Purkinje cells

Based on the strength of DE burden and evidence for altered synaptic and morphological gene signatures, we went on to characterize morphological and functional properties of PCs. We first examined the morphology of individual PCs (WT = 15 cells from 15 mice; *Chd8^+/5bpde^*^l^ = 13 cells from 13 mice; both sexes) reconstructed from WT and *Chd8^+/5bpdel^*mice at P12 (**Figs. 5A, B1-2,** see **Supplemental Table 7** for details on statistics for PC morphology and electrophysiology). Soma volume was significantly increased in *Chd8^+/5bpdel^* PCs compared with WT controls (t_26_ = 2.122, p = 0.044), indicating somatic enlargement (**Fig. 5C1**). To assess dendritic architecture, we quantified the total number of dendritic branches and total branch length (WT = 21 cells from 21 mice; *Chd8^+/5bpde^*^l^ = 13 cells from 13 mice; both sexes). Neither the number of branches (t_32_ = 1.623, p = 0.114) nor total branch length (t_32_ = 0.430, p = 0.670) differed significantly between genotypes (**Figs. 5C2-3**). Finally, we analyzed branch distribution as a function of distance from the soma (**Fig. 5D**). Mixed-effects ANOVA analysis revealed a significant effect of distance from the soma (F_(40, 800)_ = 48.05, p < 0.001) and a significant distance-by-genotype interaction (F_(3.17, 37.5)_ = 4.652, p = 0.007), whereas the main effect of genotype was not significant (F_(1, 20)_ = 0.006, p = 0.940). These findings indicate that, although the overall extent of dendritic branching was not altered, the spatial distribution of branches across the dendritic arbor differed between WT and *Chd8^+/5bpdel^* PCs, with mutant neurons exhibiting increased proximal dendritic branching within 16-52 μm of the soma.

**Figure 5:**
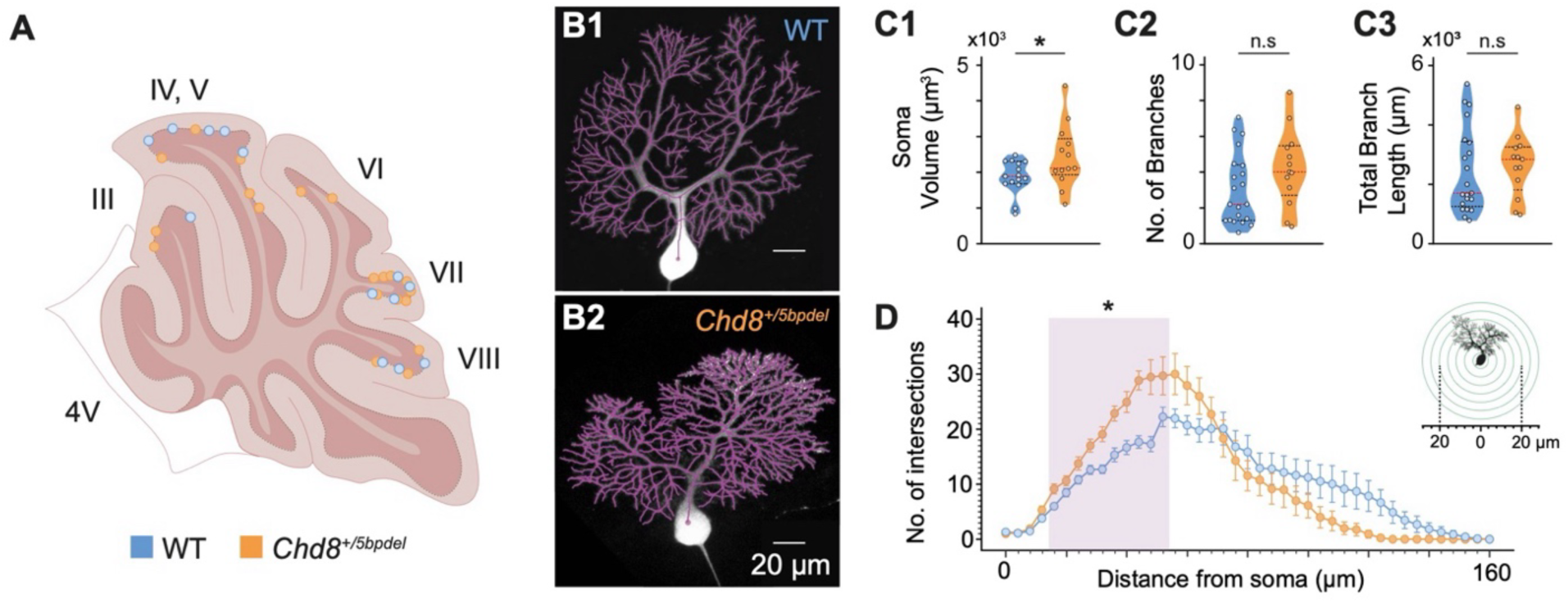
Morphological characterization of Purkinje cells in WT and *Chd8^+/5bpdel^* mice. **A**) Schematic of sagittal CB section with demarcated location of biocytin-filled PCs included in the morphological analyses; blue dots: WT; orange dots: *Chd8^+/5bpdel^*. **B1-2**) Example images of biocytin-filled WT and *Chd8^+/5bpdel^* PCs, overlaid with dendritic reconstructions (purple tracings) used for Sholl analyses. **C1-3**) Quantification of PC soma volume (**C1**), total number of branches (**C2**), and total dendritic length (**C3**). In violin plots, red dashed line: median; black dashed lines: first and third quartiles. **D**) Sholl analysis of dendritic intersections as a function of distance from PC soma, quantified using concentric circles spaced 4 μm apart (inset). Shaded area denotes significant difference between WT and *Chd8^+/5bpdel^* using a mixed-effects ANOVA followed by a false discovery rate (FDR)-corrected post hoc analysis. * denotes p-value < 0.05.

Next, we examined electrophysiological phenotypes of PCs in acute CB slices from P12 mice using whole-cell patch clamp electrophysiology. To evaluate circuit activity, we measured spontaneous PC firing from WT and *Chd8^+/5bpdel^*mice (WT = 26 cells from 7 mice; *Chd8^+/5bpde^*^l^ = 15 cells from 5 mice; both sexes) (**Fig. 6A1**). *Chd8^+/5bpde^*^l^ PCs displayed a trend toward higher spontaneous firing rates compared with WT, however, this difference did not reach statistical significance (t_39_ = 1.845, p = 0.073) (**Fig. 6A2**). Similarly, the coefficient of variation (CV) of ISI was not significantly different between genotypes (t_39_ = 0.563, p = 0.576) (**Fig. 6A3**). In contrast, CV2 was significantly increased in *Chd8^+/5bpdel^*PCs (t_39_ = 2.078, p = 0.044), indicating increased local variability in spike timing and reduced firing regularity (**Fig. 6A4**). Next, we assessed passive membrane properties of PCs (WT = 28 cells from 7 mice; *Chd8^+/5bpdel^*= 22 cells from 5 mice; both sexes) **(Figs. 6B1-4**). Input resistance (R_input_) was significantly reduced in *Chd8^+/5bpdel^* PCs (t_48_ = 2.271, p = 0.028) (**Fig. 6B2**), and membrane time constant (τ) was significantly increased (t_48_ = 2.058, p = 0.045) (**Fig. 6B3**). Membrane capacitance (C_m_) was also elevated in mutant PCs (t_48_ = 2.689, p = 0.010) (**Fig. 6B4**). No differences were detected in sag ratio between genotypes (t_48_ = 0.433, p = 0.667).

**Figure 6:**
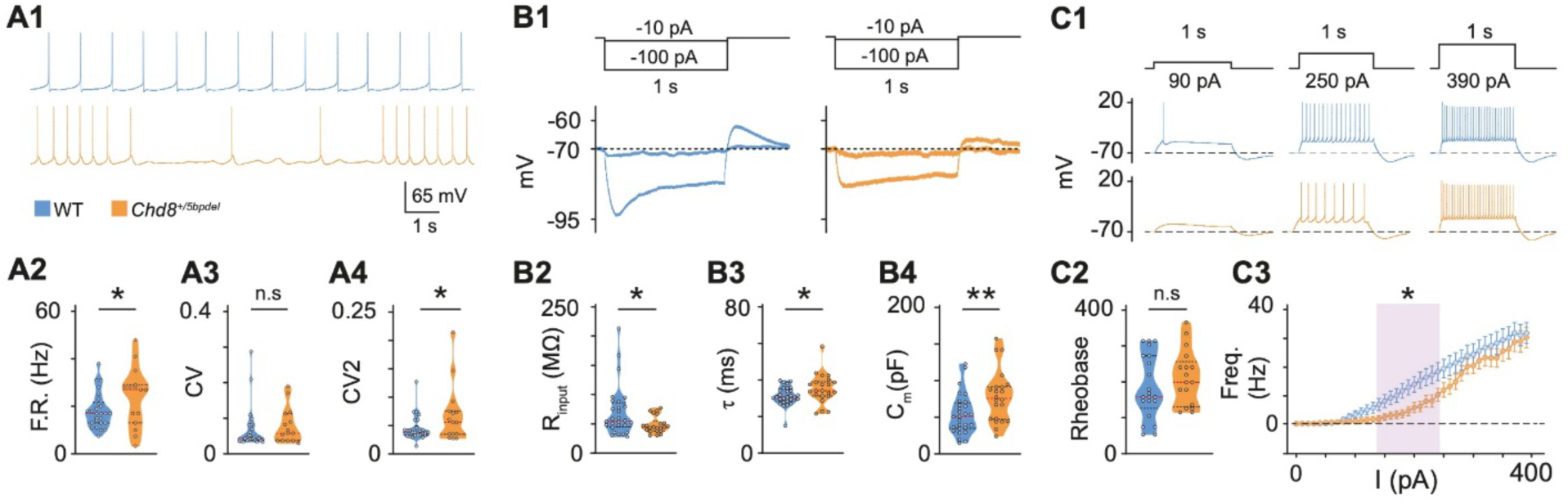
Intrinsic firing and membrane properties of Purkinje cells in WT and *Chd8^+/5bpdel^* mice. **A1**) Example spontaneous firing rate traces from WT and *Chd8^+/5bpdel^* PCs. **A2-4**) Violin plots of spontaneous firing rate (**A2**), coefficient of variation (CV; **A3**), local coefficient of variation (CV2; **A4**) for WT and *Chd8^+/5bpdel^*. **B1**) Representative voltage responses to hyperpolarizing current injections. **B2-4**) Violin plots of input resistance (R_input_; **B2**), membrane time constant (τ) **B3**), and membrane capacitance (C_m_, **B4**) for WT and *Chd8^+/5bpdel^*. **C1)** Representative voltage responses to depolarizing current injections. **C2**) Rheobase. **C3**) Frequency-current (F-I) curves. Shaded area denotes significant difference between WT and *Chd8^+/5bpdel^* using mixed-effects ANOVA followed by false discovery rate (FDR)-corrected post hoc analysis. All panels: In violin plots, red dashed line: median; black dashed lines: first and third quartiles. Comparisons between two independent groups used two-tailed unpaired *t*-tests, except for C3. *p-value < 0.05, n.s.: p-value > 0.05.

For active properties, we analyzed single action potential (AP) waveform properties and found no significant differences between genotypes in threshold (t_46_ = 1.346, p = 0.185), amplitude (U = 198, p = 0.076), half-width (U = 225, p = 0.228), rise time (t_46_ = 0.064, p = 0.949), decay time (t_46_ = 1.846, p = 0.071), rise slope (t_46_ = 1.048, p = 0.300), decay slope (t_46_ = 1.493, p = 0.142), fast afterhyperpolarization (t_46_ = 1.022, p = 0.312) or medium afterhyperpolarization (t_46_ = 1.684, p = 0.099) (**Supplemental Fig. 9 and Supplemental Table 7**). Likewise, spike frequency adaptation was unaltered (SFA ratio: U = 179, p = 0.193). We further characterized intrinsic excitability (WT = 27 cells from 7 mice, both sexes; *Chd8^+/5bpdel^* = 21 cells from 5 mice, both sexes), using a series of depolarizing current injections (**Figs. C1-3**). Although rheobase was not significantly different between WT and *Chd8^+/5bpdel^* PCs (t_36_ = 0.977, p = 0.335; WT = 21 cells from 7 mice; *Chd8^+/5bpdel^* = 17 cells from 5 mice) (**Fig. 6C2**), mutant neurons exhibited a slight trend toward higher values. We therefore probed the frequency-current (F-I) relationship across the full range of injected currents. This analysis revealed a significant effect of current injection (F_(1.84, 84.56)_ = 77.62, p < 0.0001) and a significant current-by-genotype interaction (F_(39, 1794)_ = 1.560, p = 0.015), whereas the main effect of genotype was not significant (F_(1, 46)_ = 2.279, p = 0.138). Post-hoc multiple-comparison analysis identified significant differences between genotypes across a range of intermediate current injections (140-240 pA; **Fig. 6C3**). Within this range, *Chd8^+/5bpdel^* PCs generated fewer action potentials than WT neurons, resulting in a rightward shift of the F-I relationship. However, firing frequencies progressively converged at higher current injections, indicating that maximal firing output was largely preserved.

To further characterize the input-output relationship, F-I curves were fitted with a sigmoidal function and neuronal gain was estimated from the fitted responses. *Chd8^+/5bpdel^* PCs exhibited a steeper gain than WT neurons (145.43 vs.114.75 Hz/nA), consistent with a more abrupt increase in firing frequency once action potential generation was initiated. Together, these findings suggest that *Chd8* haploinsufficiency could alter recruitment of PCs in response to depolarizing inputs, whereas the ability to achieve normal firing rates upon strong stimulation is maintained.

To assess effects on synaptic input associated with *Chd8* haploinsufficiency, miniature excitatory and inhibitory postsynaptic currents (mEPSCs and mIPSCs) were recorded from PCs (mEPSC: WT = 18 cells from 7 mice; *Chd8^+/5bpde^*^l^ = 14 cells from 6 mice; mIPSC: WT = 34 cells from 18 mice; *Chd8^+/5bpde^*^l^ = 37 cells from 14 mice; both sexes) (**Figs. 7A,B1-2**). Analysis of mEPSCs revealed a significant increase in event frequency in *Chd8^+/5bpde^*^l^ PCs (t_30_ = 2.130, p = 0.042; **Figs. 7C1-2**), suggesting enhanced excitatory synaptic input. In contrast, mEPSC amplitude was not significantly different between genotypes (t_30_ = 0.429, p = 0.671; **Fig. 7C3**), indicating that postsynaptic response magnitude remained unchanged. We next examined inhibitory synaptic transmission (**Fig. 7D1**). Neither mIPSC frequency (t_69_ = 1.222, p = 0.226) (**Fig. 7D2**) nor mIPSC amplitude (t_69_ = 0.265, p = 0.792) (**Fig. 7D3**) differed significantly between WT and *Chd8^+/5bpde^*^l^ PCs, suggesting that inhibitory synaptic transmission is largely preserved in mutant animals. To further evaluate genotype-dependent changes in synaptic event properties, cumulative probability distributions were compared using two-sample Kolmogorov–Smirnov tests (**Figs. 7C4,D4**). The distribution of excitatory events differed significantly between WT and *Chd8^+/5bpdel^*mice (D = 0.298, p < 0.001), indicating a substantial shift in the overall distribution of mEPSCs. In contrast, inhibitory event distributions did not differ significantly between genotypes (D = 0.030, p = 0.622). These findings indicate a selective increase in excitatory, but not inhibitory, synaptic transmission onto *Chd8^+/5bpde^*^l^ PCs.

**Figure 7:**
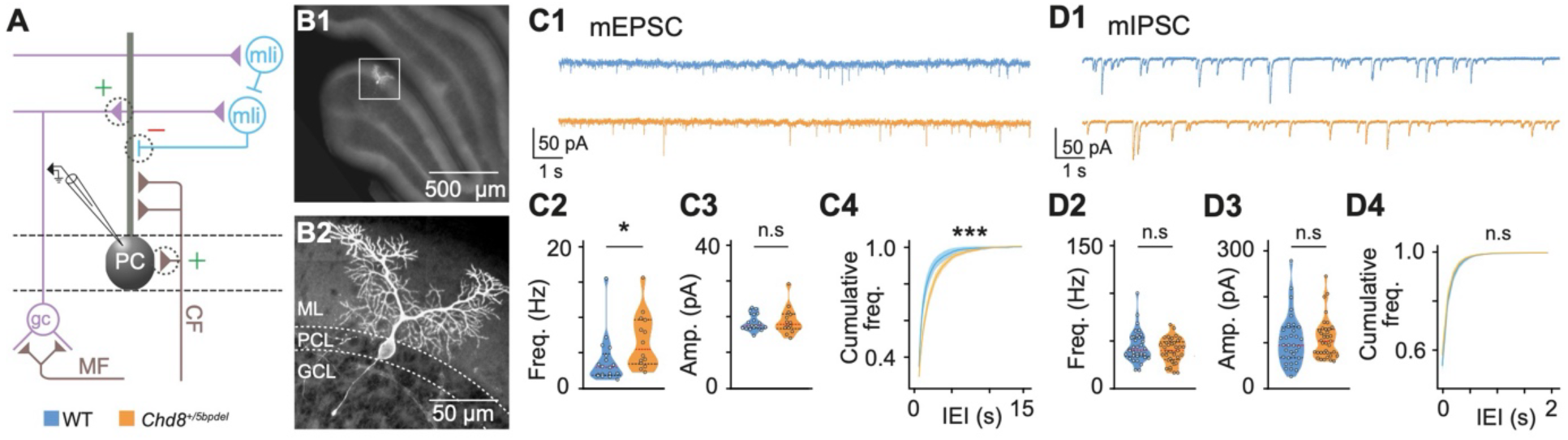
Miniature synaptic transmission in Purkinje cells from WT and *Chd8^+/5bpdel^* mice. **A**) Simplified schematic of CB cortical circuitry. MLI: molecular layer interneuron; MF: mossy fiber; CF: climbing fiber; PC: Purkinje cell; gc: granule cell. ML, molecular layer; PCL, Purkinje cell layer; GCL: granule cell layer. **B1**) Representative image of a recorded PC filled with biocytin. **B2**) Zoom in. **C1**) Example miniature excitatory postsynaptic current (mEPSC) traces recorded from WT and *Chd8^+/5bpdel^* PCs. **C2-4**) Frequency (**C2**), amplitude (**C3**), and cumulative distributions of the inter-event interval (IEI; **C4**) of mEPSCs from WT and *Chd8^+/5bpdel^*PCs. **D1**) Representative miniature inhibitory postsynaptic current (mIPSC) traces. **D2-4**) Frequency (**D2**), amplitude (**D3**), and cumulative distributions of IEI (**D4**) of mIPSCs for the two genotypes. All panels: in violin plots, red dashed line: median; black dashed lines: first and third quartiles. Comparisons between two independent groups were performed using two-tailed unpaired *t*-tests, and cumulative distributions were compared using the two-sample Kolmogorov-Smirnov test. *p-value < 0.05, n.s.: p-value > 0.05.

Collectively, our morphological and electrophysiological analyses (**Supplemental Table 9**) highlight a striking correlation between structural expansion and altered functional excitability in *Chd8*^+/*5bpdel*^ PCs. The morphological increases in soma volume and proximal dendritic branching align tightly with the higher membrane capacitance and decreased input resistance measured in our recordings. Such an increase in cell membrane would act as a capacitive load, potentially explaining the requirement for larger depolarizing inputs to initiate firing in mutant PCs. Concurrently, the extra surface area provided by proximal dendritic expansion could facilitate the formation of additional excitatory synapses, though the elevated mEPSC frequency might also represent a compensatory physiological response to the reduced PC intrinsic excitability. Ultimately, the selective enhancement of excitatory input is consistent with the transcriptomic upregulation of glutamatergic pathways, collectively pointing to PCs as a central locus of circuit dysfunction in the developing *Chd8^+/5bpde^*^l^ CB.

### Global transcriptomic dysregulation persists into adulthood in the *Chd8^+/5bpdel^* cerebellum

To determine whether early *Chd8*-dependent neurodevelopmental perturbations resolve over time or translate into enduring network pathology, we profiled the adult CB transcriptome through matched bulk RNA-seq at both P12 and P60. We analyzed whole CB dissections from P12 WT (N = 11) and *Chd8^+/5bpdel^* (N = 11) mice, as well as P60 WT (N = 16) and *Chd8^+/5bpde^*^l^ (N = 16) mice (both sexes for all groups). Principal component analysis (PCA) didn’t separate samples by genotype in PC1 or PC2 (**Supplemental Fig. 10A,B).** Bulk DEGs were identified using DESeq2, with PC1, PC2, and sex included as covariates (**Supplemental Fig. 10C)**. At P12, there were 833 downregulated genes (p< 0.05), 42 of which passed FDR < 0.1, and 778 upregulated genes (p < 0.05), 59 of which passed FDR < 0.1 (**Fig. 8A**). At P60, there were 1,727 downregulated genes (p < 0.05) 448 of which passed FDR < 0.1 and 1,200 upregulated genes (p < 0.05), 254 passing FDR < 0.1 (**Fig. 8B**). See **Supplemental Table 8** for full DE results. At both ages, there was high correlation in DEG effects between males and females (**Supplemental Fig. 10D,E**). *Chd8* was the most significant DEG at both ages (**Fig. 8C**), confirming *Chd8* haploinsufficiency in *Chd8^+/5bpdel^* CB. Eighty-two genes passed FDR < 0.1 at both ages, including *Tbr1*, a marker of PNs in DCN, and *Ucp2*, a mitochondrial regulator (**Fig. 8C**). DEG effects showed strong directional correlation between the two ages (**Fig. 8D**). P12 DEGs (FDR < 0.1) were largely expressed across cerebellar cell types, (**Supplemental Fig. 10F)**. Together, these results indicate that transcriptomic signatures detectable in bulk at P12 persist into adulthood, with evidence for increased effect size and significance in the adult CB.

**Figure 8:**
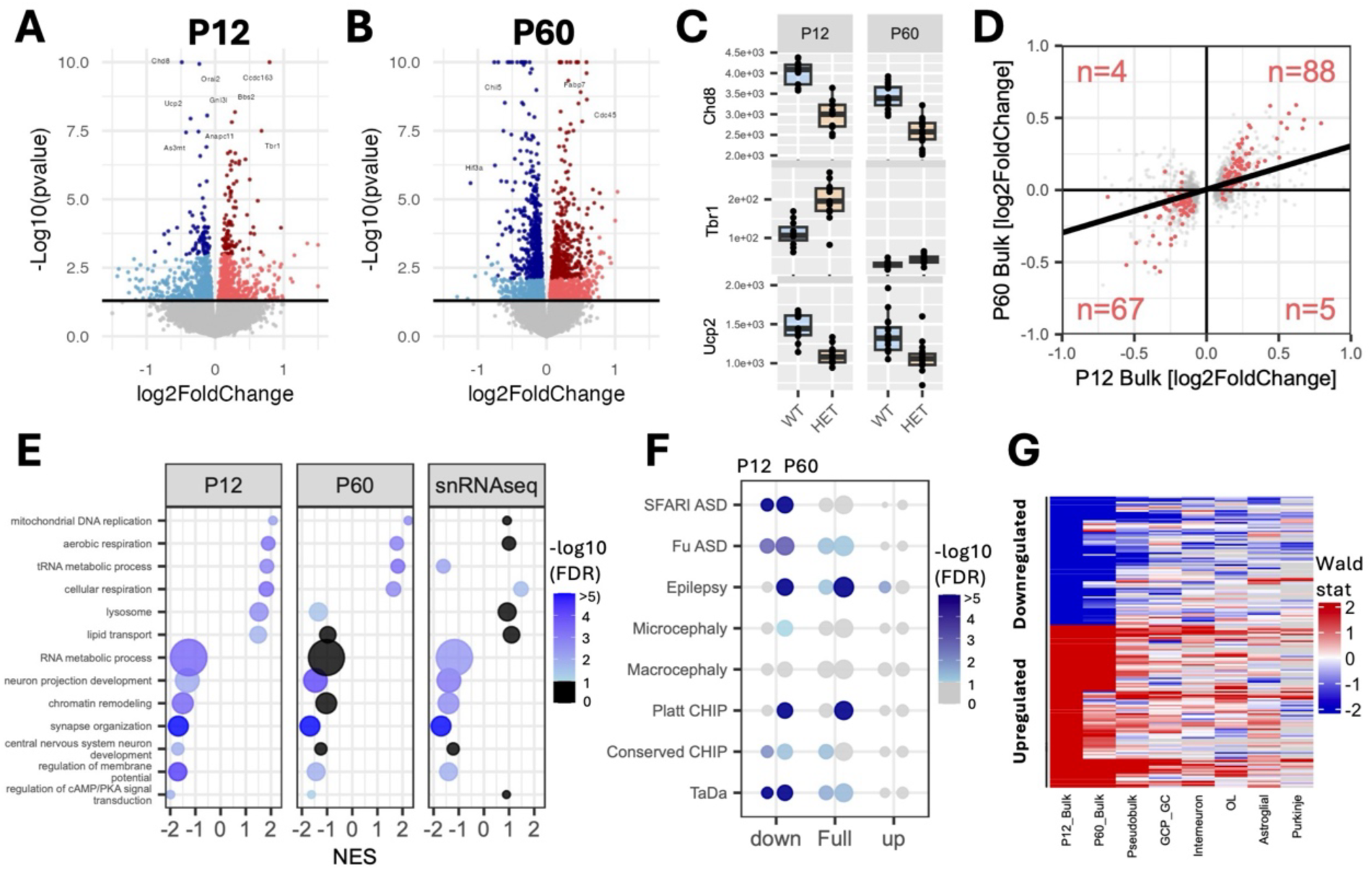
Developmental transcriptomic dysregulation persists into adulthood. **A**) Volcano plot showing DE of P12 CB bulk RNA-seq data between wildtype and *Chd8^+/5bpdel^* samples. Color indicates significance and DE; light red (upregulated, p < 0.05), dark red (upregulated, FDR < 0.1), light blue (downregulated, p < 0.05), dark blue (downregulated, FDR < 0.1), p > 0.05 (grey). **B**) Volcano plot showing differential expression of P60 CB bulk RNA-seq data between wildtype and *Chd8^+/5bpdel^* samples. Color scheme as in A). **C**) Boxplots showing normalized expression by genotypes for *Chd8* and other top representative DEGs (FDR < 0.05) across both age groups. **D**) Scatterplot comparing effect sizes (log2 fold change) of DEGs between P12 and P60 bulk RNA-seq data. Dot color denotes significance; grey (p < 0.05) and red (FDR < 0.1) with the number indicated the number of FDR genes in each quadrant. Correlation between P12 FDR genes derived from a linear model, R^2^ = 0.67. **E**) Dot plot showing a curated set of significant (FDR < 0.05) GO terms from the P12 bulk RNA-seq in the P60 bulk RNA-seq and P12 snRNA-seq pseudobulk data. Upregulated (NES > 0) and down-regulated (NES < 0) gene sets from GSEA ranked by DESeq2 Wald test. Color indicates the signed enrichment of the gene set over background and size indicated size of enriched gene set. **F**) Dot plot showing the results of permutation tests of DEGs (p < 0.05) in the bulk RNA-seq at P12 and P60 against disease-associated and curated *Chd8*-associated gene sets. Size indicates enrichment of the gene set over background and color the strength of significance. **G**) Heatmap showing the differential expression of P12 DEGs (FDR < 0.1) in the P60 bulk RNA-seq, snRNA-seq pseudobulk (whole dataset), and snRNA-seq pseudobulk (within cell type). Color denotes DESeq2 Wald test score, empty cells did not pass counts criteria for DE testing in snRNA-seq data.

We used GSEA to test whether perturbations to functional processes were similarly conserved between ages (**Fig. 8E, Supplemental Table 9**). Among the most significantly upregulated GO terms at P12 were *mitochondrial DNA replication* and *aerobic respiration*, both of which were also significantly upregulated in the P60 data. Downregulated processes such as *synapse organization* and *neuron projection development* were likewise significant at both timepoints. Basal cellular process dysregulation, including RNA metabolic process and chromatin remodeling, shared concordant effect direction between ages, but did not reach significance in the adults. We also tested concordance between the bulk RNA-seq results and pseudobulk DE calculated from the combined cells of all cell types in the snRNA-seq data. The majority of effects were conserved between bulk and snRNA-seq. Notably, the decreased neuronal and synaptic signatures observed in the scRNA-seq pseudobulk appear to be driven by a reduced number of PNs, likely including those originating from the DCN. We used overrepresentation analysis to test whether *Chd8*-associated and other disease-relevant gene sets were perturbed in our bulk RNA-seq data (**Fig. 8F**). We found conserved downregulation of ASD-associated genes and *Chd8* ChIP-seq targets at both timepoints, whereas epilepsy-associated genes showed strong, adult specific enrichment for upregulated DEGs. Finally, comparison of bulk and snRNA-seq data at the gene level showed trends of concordant effects but did not reach significance (**Fig. 8G**).

Together, the comparison of P12 and adult bulk RNA-seq demonstrates that CB transcriptional consequences of *Chd8* haploinsufficiency persist from postnatal development into adulthood and become more extensive by P60. Shared alterations included increased mitochondrial and aerobic respiration programs, reduced synaptic and neuronal-development programs, and conserved downregulation of ASD-associated genes and CHD8 targets. Concordance between bulk and single-nucleus RNA-seq further indicates that these effects are robust across transcriptomic preparations, with bulk RNA-seq identifying general changes and snRNA-seq resolving cell-type-specific context.

## Discussion

Here we characterized effects of disease-relevant *Chd8* haploinsufficiency on the developing mouse CB via an integrated transcriptomic, morphological and functional approach. Our experiments revealed molecular, developmental, morphological, and physiological abnormalities that were not readily appreciated from evaluation of gross anatomy. Major findings included shifts toward more mature transcriptional states across developing cell lineages, shared and cell-type-specific transcriptional pathology, pronounced molecular and cellular vulnerability of PCs, and persisting transcriptomic dysregulation into adulthood. These results greatly extend previous work on *Chd8* in the CB, demonstrating that germline constitutive heterozygous loss-of-function mutation is sufficient to disrupt CB development and function. Broadly, our findings illustrate how NDD-linked single-copy loss-of-function mutations in chromatin remodelers such as *Chd8* can produce widespread, cell-type-specific pathology within brain regions that are understudied, despite their critical contribution to cognitive and social behavior, and further point to the CB as a potential contributor to CHD8-NDD pathology.

Our overall objective was to determine whether germline *Chd8* haploinsufficiency alters CB development. In contrast to the severe hypoplasia and disrupted lamination caused by homozygous deletion of *Chd8* in granule cell precursors^25,26^, the heterozygous germline loss-of-function we studied here did not impact overall volume, laminar organization, or the abundance of major cell classes in the P12 cerebellar cortex. Nevertheless, GC, MLI, and OL contained fewer immature cells and proportionally more mature states. Pseudotime and developmental gene set analyses similarly indicated more advanced transcriptional states, including increased neuronal synaptic programs and oligodendrocyte myelination programs. These results suggest that reduced *Chd8* dosage in the developing CB alters the timing and/or coordination of postnatal CB maturation. Shifts in the maturation of cell types have been observed in other *Chd8* mouse models and organoid studies^23,48,49^, but this is the first examination within the disease-relevant *Chd8* haploinsufficient CB and its cell types. Mistimed development of CB cell types could disrupt the canonical assembly and maturation of CB circuitry and downstream targets including thalamus and prefrontal cortex^50–52^, leading to lasting dysfunction.

Our second objective was to identify the CB cell types most susceptible to reduced *Chd8* dosage. Differential expression, cell prioritization, gene set enrichment, and co-expression network analyses converged on PCs as the most affected CB population at P12. PCs exhibited the greatest DE burden and were the only cell type showing significant enrichment of ASD-associated genes, all despite the relatively limited PC representation in the snRNA-seq datasets. RNAscope validation of the increases in *Corin* expression and proportion of *Corin*-positive PCs corroborates the cell-type-specific changes detected by snRNA-seq and raises the possibility that *Chd8* haploinsufficiency alters *Corin*-associated PC state or subtype identity. Prior work focusing on conditional PC knockout mice did not identify substantial phenotypes in either homozygous or heterozygous mice^26^. However, the CRE driver used in that study does not drive recombination until later in PC development^53^. Thus, the phenotypes we observed at the molecular and functional level would have been missed if associated with early *Chd8* requirement in PC specification.

Our combined single-nucleus and functional analyses suggest that *Chd8* haploinsufficiency drives a network-wide, maladaptive acceleration of CB circuit maturation. Our trajectory analyses reveal that both excitatory GCs and inhibitory MLIs prematurely shift toward more mature states at P12. However, the functional consequences of this acceleration differ by cell type. GCs robustly upregulate presynaptic neurotransmitter secretion pathways, which could account for the increased excitatory synaptic drive observed in downstream PCs. In contrast, maturing MLIs preferentially upregulate postsynaptic machinery to receive inputs, providing a transcriptomic explanation for why we observed no corresponding increase in inhibitory transmission onto PCs. At the center of this network, PCs physically remodel, enlarging their somata and expanding complexity of their proximal dendrites, perhaps to accommodate this premature synaptic integration. The structural expansion would be expected to create a larger sink of synaptic current in PCs and could explain the requirement for more depolarizing input to fire action potentials. Ultimately, the combination of altered intrinsic gene expression and the increased excitatory synaptic drive could disrupt the stability of PC baseline firing, as revealed by changes in CV2. This working model could potentially explain how *Chd8* mutations, by triggering asynchronous developmental acceleration across distinct cell lineages, culminate in a functionally mistuned CB circuit. Our results echo work in other ASD/NDD mouse lines identifying PCs as a critical axis of pathology^9,10^. Further work will be required to determine the relevance of phenotypes identified here for CB-relevant behaviors.

A final objective was to determine whether CB transcriptional pathology persists in adulthood. Matched bulk RNA-seq of P12 and P60 CB tissue revealed a strong directional concordance in transcriptional effects, indicating that many early molecular abnormalities are not resolved during maturation. Specifically, we observed sustained upregulation of mitochondrial and aerobic respiration programs alongside a conserved downregulation of ASD-associated genes and *Chd8* targets across both ages. Bulk sequencing also showed a shared downregulation of synaptic and neuronal development gene sets, and our single-nucleus data suggest this could be driven by a lasting reduction in PN populations, a finding consistent with our previous report of reduced DCN volume^21^. Notably, an enrichment of upregulated epilepsy-associated genes emerged specifically in the adult CB. This adult-specific signature aligns with partially penetrant clinical comorbidity of seizures in CHD8-NDD^17,19^ and suggests that early developmental miswiring in the CB may contribute to altered susceptibility to network hyperexcitability in adulthood. Many of these altered functional programs, particularly aerobic respiration and RNA processing, parallel findings from *Chd8* mouse studies focused on the cerebral cortex^21,54^. These conserved transcriptomic signatures suggest that *Chd8* haploinsufficiency exerts shared, systemic impacts across different brain regions, providing a compelling foundation for future functional follow-up.

Several limitations should be considered when interpreting these findings. P12 represents a dynamic period of CB development during which neuronal differentiation, circuit assembly, and synaptogenesis are ongoing^29,55^. At least some of the observed transcriptional and physiological phenotypes may reflect shifts in developmental timing rather than stable pathological states. Also, whether the observed changes in PC excitatory transmission, intrinsic excitability, and dendritic organization persist in adulthood remains to be investigated. In addition, the snRNA-seq was not sufficiently powered to identify sex-specific effects, which have been reported in other *Chd8* mouse studies^56^, though we did not see strong sex effects in exploratory snRNA-seq analysis or in our bulk datasets, and we also did not detect sex differences in preliminary electrophysiology. Moreover, snRNA-seq measures steady-state nuclear RNA and therefore does not fully capture whole-cell transcript abundance or stability. The reduced sampling after subdivision into cell types limits the detection of modest or low abundance expression changes, even with pseudobulk analysis. Bulk RNA-seq offers greater sensitivity but cannot resolve cell-type-specific effects and may be influenced by genotype-associated or technical differences in cellular composition. Finally, we did not extend our work to behavioral assays in this study. We and others have previously established behavioral phenotypes in *Chd8* haploinsufficient mice^21–23,54,56–58^. Crucially, because our model utilizes a constitutive mutation, behavioral outputs reflect whole-brain pathology and cannot be conclusively attributed to the specific CB microcircuit abnormalities identified here.

In conclusion, our results demonstrate that *Chd8* haploinsufficiency disrupts CB development and neuronal function. Future work will define the developmental onset of these phenotypes, determine whether PC alterations are cell-autonomous and persist into mature developmental ages, and establish how altered cell maturation, myelination, and excitatory input interact to shape PC output. Regardless, the convergence of developmental state shifts across multiple lineages and the pronounced molecular, morphological, and physiological changes in PCs highlights the CB as a locus of early *Chd8*-associated pathology.

## Acknowledgements

This work was supported by the National Institute of Mental Health (NIMH) grants R21 MH126413 (to A.S.N. and D.F.) and R21 MH132834 (to D.F. and A.S.N.); R01 MH120513 (to A.S.N.); T32 MH073124 (through M.I.N.D Institute, to C.P.C); F31 MH119789 (to N.S); F31 DA062491 (to E.F.); NIH F31 predoctoral fellowships F31 HD113328 (to S.L.); NIMH Ruth L. Kirschstein National Research Service Award (NRSA) Institutional Research Training Grants T32 MH082174 (Basic Neuroscience, supporting E.F. and A.D.A.) and T32 MH112507 (Learning, Memory, and Plasticity, supporting A.D.A. and N.S.).

## Author Contributions

N.S., E.M., C.P.C., D.F., and A.S.N. conceptualized the study. N.S., E.M., A.D.A., S.J.J., J.L.D., S.L., K.J., D.R., M.C., C.A. and C.P.C performed experiments. E.F., N.C., K.C., and E.G.A. contributed computational tools. N.S., E.M., A.D.A., J.L.D., and D.F. analyzed data. N.S., E.M., S.J.J., C.P.C., D.F., and A.S.N. wrote the manuscript. N.S., E.F., S.L., C.P.C., D.F., and A.S.N. procured funding that supported the research.

## Materials and Methods

### Animals

Generation and characterization of Cas9-mediated 5bp frameshift deletion in exon 5 of *Chd8* in mice (here referred to as *Chd8^+/5bpdel^* mice) was previously described^21^. All protocols utilized in the generation of mouse CB samples were approved by the Institutional Animal Care and Use Committees (IACUC) at the University of California Davis. Mice were housed in a temperature-controlled vivarium maintained on a 12-h light-dark cycle. We made efforts to minimize pain, distress and the number of animals used in the study.

### Histology and Immunohistochemistry

IHC was performed on free-floating serial sections from P12 mouse CB according to our established protocols. Anesthetized mice were perfused transcardially with 4% (w/v) paraformaldehyde (PFA; EMS Diasum, Hatfield, PA, USA) in phosphate buffered saline (PBS; Sigma-Aldrich, St. Louis, MO, USA). Brains were post-fixed in 4% paraformaldehyde, cryoprotected in 30% sucrose in PBS, and sectioned at 40 µm using a cryostat. Sections were permeabilized (PBS-0.5% Triton X-100, 20 min at RT) and non-specific binding was blocked (5% skimmed milk dissolved in 0.1% PBST, 1 h at RT or overnight at 4 °C). Slices were incubated overnight with primary antibody (anti-Calb1, PA5-50366, ThermoFisher Scientific, 1:50), then AlexaFluor-conjugated secondary antibodies (1:1000, for 1 hr at RT), and mounted on slides with ProLong Gold Antifade Mountant (Invitrogen).

For CB volume measurements, mice were anesthetized with the anesthetic cocktail and perfused transcardially with 4% (w/v) PFA. Brains were post-fixed in 4% PFA for 24 h and transferred to 30% sucrose in PBS for overnight incubation at 4°C. Brains were coronally sectioned (80 μm) using a cryostat (3050S, Leica Microsystems, Wetzlar, Germany), stained with DAPI (1:20,000; Thermo Fisher Scientific, Waltham, MA, USA), mounted onto glass slides, and cover-slipped with a Mowiol-based antifade glycerol solution.

For PC monolayer analysis, P12 sagittal sections were stained with an anti-CALB1 antibody. High-power microscopy using a Keyence epifluorescence microscope was used to qualitatively define the organization of PCs within lobule simplex of the CB. For each sample, we analyzed 3 areas per lobule producing 18 regions of interest (ROI) per condition. For each ROI, the monolayer or multilayer organization of PCs was determined visually by 3 independent raters that were blind to genotype.

To qualitatively define the expression of CHD8 in the P12 CB, serial sagittal sections were co-stained with an anti-CALB1 antibody and anti-CHD8 C-terminus antibody (Fortis Life Sciences, A301-225A). CHD8 expression was defined across the ML, PCL, GCL, and DCN.

Fluorescence images were acquired using a Keyence epifluorescence microscope (BZ-X1000, Keyence Corporation, Osaka, Japan) or a Zeiss LSM800 confocal microscope equipped with Airyscan (60x, Carl Zeiss AG, Oberkochen, Germany). Maximum intensity projections of confocal z-stacks were generated using ZEN software (Carl Zeiss AG, Oberkochen, Germany) or ImageJ and used for subsequent morphological analyses.

### Cerebellar dissections for transcriptomic experiments

*Chd8^+/5bpdel^* and WT littermate mice were anesthetized with isoflurane and euthanized by cervical dislocation. For snRNA-seq P12 tissue, whole brains were extracted and ∼450 µm coronal slices containing the CB were prepared using a sectioning mold (Civic Instruments #5325). Slices spanning approximately –2.5 mm to –3.0 mm relative to lambda were collected, using coordinates from the Paxinos mouse brain atlas as an approximate anatomical reference analogous to corresponding adult landmarks, rather than as precise stereotaxic coordinates for this age group. Extracerebellar tissue contained within the boundaries of each coronal slice was retained to maintain dissection consistency across samples. For developmental (P12) and adult (P60) bulk RNA-seq, tissue was collected by dissecting one half of the whole CB, with the left or right hemisphere selected randomly across samples. All dissected tissue was flash-frozen in dry ice and stored at –80°C until processing.

### General bioinformatics

Bioinformatic analysis was performed using R programming language (version 4.4.2) (R Development Core Team, 2015) and RStudio integrated development environment (version 2026.1.2.418) (Team R, 2018). Plots were generated using ggplot2 R package (version 3.4.0) and pheatmap (version 1.0.13).

## Single-nucleus RNA-seq

### Nuclei isolation for snRNA-seq and pooling strategy

Nuclei were prepared using two methods, a modified version of the Frankenstein nuclei isolation protocol^59^ used for the initial samples, and the 10X Genomics nuclei isolation kit for latter samples. For the first method, frozen tissue was processed into nuclei using 2mL glass dounce homogenizers containing 2mL of EZ Prep lysis buffer. Each solution was transferred to a 15 mL conical tube, then an additional 6 mL of EZ Prep lysis buffer was added to each conical tube and the tubes were incubated on ice for 5 min. Homogenates were centrifuged at 500 x g for 5 minutes at 4°C, the supernatant was discarded, and the pellet was washed twice in NSB (1X PBS with 0.01% BSA and 0.1% RNAse inhibitor). The final pellet was resuspended in 1mL NSB then passed through a 20 μM cell strainer. Individual nuclei preps were counted and integrity was assessed with a Countess cell counter, using a DAPI nuclear stain. For the 10X Genomics nuclei isolation kit, we used to isolate nuclei for the final 4 samples following manufacturer protocol. Final nuclei counts were determined using a LUNA-FL cell counter and Acridine Orange/ Propidium Iodide stain. For all samples, each run included equal amounts of one male and one female replicate of opposite genotype. Downstream genotyping in the snRNA-seq data found one genotyping error, with one pool was determined to have only *Chd8^+/5bpdel^*samples of both sexes, and one sample with two females, with correct genotype and sex used for downstream analysis. Nuclei suspensions had concentrations adjusted to target 10,000 nuclei for sequencing per pool, then used as input for the 10X Genomics 3’ Gene Expression Assay. Libraries were generated according to the v3 protocol by the UC Davis DNA Technologies Core. For the first 4 pools, libraries were sequenced with the Illumina NovaSeq 6000 at 80,000 reads per nucleus. The remaining 3 pools were sequenced with the Element Biosciences AVITI at 80,000 reads per nucleus. Raw fastq files and integrated Seurat object available on GEO.

### snRNA-seq bioinformatics

#### Pre-processing and initial QC

De-multiplexed sequencing data were obtained from the UC Davis DNA Technologies Core as FASTQ files. FASTQ files were passed through the Cell Ranger (10X Genomics) *count* pipeline for alignment to the mouse mm10 genome, filtering, and creation of feature-barcode matrices. Ambient RNA removal was performed using SoupX^60^ (version 1.6.2), and Seurat objects were generated using Seurat^45^ (version 5.4.0). These Seurat objects were subjected to initial clustering, and each resulting cluster was filtered based on cluster-specific thresholds for UMI count, feature count, and percentage of mitochondrial reads, which were determined using standard deviation criteria. For doublet removal, the scDblFinder^61^ package (version 1.20.2) was used to assign nuclei as ‘Singlet’ or ‘Doublet’. Predicted doublets and clusters primarily composed of doublets were removed before proceeding with downstream analyses.

#### Assigning sex and genotype

Nuclei were assigned as ‘male’ or ‘female’ based on the expression of sex-linked marker genes located on the X and Y chromosomes. Cells were only used for differential analysis if they contained either only male or female sex-linked genes. *Chd8* genotype was assigned based on the predicted sex and pool-specific metadata. Because each pool consisted of only one male and one female of opposite genotype, all female nuclei in a pool were assumed to be one genotype and all males were assumed to be the other. To confirm genotype assignment, we used the BAM files generated by cell ranger to visualize the presence of the 5bp deletion in *CHD8*.

#### Clustering analysis

Seurat objects for each of the 6 pools were individually passed through *SCTransform* to normalize counts, then integrated using *IntegrateData*. Principal Components Analysis was performed on the ‘Integrated’ assay of the integrated data to identify top variable genes, and *FindNeighbors* was used to construct a KNN graph based on their euclidean distance in PCA space. The data were then passed through *FindClusters* to group cells into clusters using the Louvain algorithm. Clusters were visualized with uniform manifold approximation and projection (UMAP).

#### Cell type annotation

To annotate the data-driven clusters, we looked at expression of known markers of cell type in the CB derived from published single-cell datasets. All mature cell types were annotated using the cell type-specific markers^30^. Oligodendrocytes were annotated using developmental markers^34^. Markers to differentiate MLI progenitors and GC progenitors were obtained from a single-cell atlas of mouse cerebellar development^62^.

#### Cell type proportion analysis

For each cell population, nuclei of each genotype were counted, then counts were converted to a proportion of the total cells in each sample. A Student’s t-test was used to assess statistical differences in the proportions of *Chd8^+/5bpdel^* and WT nuclei for each cell type. *scDist cell prioritization analysis.* Cell-type-specific transcriptional perturbations were quantified using scDist^36^ (version 1.1.5). For each major cell type, scDist estimated the distance between WT and *Chd8^+/5bpdel^* cells in high-dimensional gene expression space using a linear mixed-effects framework that accounts for variation among biological replicates. Cell types were ranked by the magnitude of the estimated genotype-associated transcriptomic shift. Statistical significance was corrected for testing across cell types using the false-discovery rate. Cell types with FDR < 0.1 were considered significantly perturbed.

#### DE analysis

A pseudo-bulk approach was used to identify DEGs in each CB cell population. Each cluster was extracted as a subset of the main Seurat object, then raw counts for genes were passed through the *DESeq2* (version 1.46.0) pipeline to compare *Chd8^+/5bpdel^*and WT nuclei within that subset. Only genes with a minimum of 10 counts in 5 samples were considered for DE analysis. PC1, PC2, and sex were included as covariates in the *DESeq2* statistical model. DEGs with p-value < 0.05 were deemed significant.

#### Burden analysis

For each cell type, DE burden was calculated as the percentage of DEGs relative to the total expressed genes that passed the minimum expression threshold to be included in the DE test. *hdWGCNA.* High-dimensional weighted gene co-expression network analysis (hdWGCNA) was applied to construct gene co-expression networks for major cell classes (Interneuron, OL, Astroglial, GCP_GC) using the hdWGCNA^44^ package (version 0.4.12). Nuclei of specific cell types were grouped by sample and passed through the *MetacellsByGroups* function to create metacells. Custom parameters for *’k’*, *’max_shared’*, and *’min_cells’* were chosen and the optimal soft power was determined using *TestSoftPowers*. The co-expression network was built with *ConstructNetwork*, and modules were detected via hierarchical clustering. Module number, size, and gene composition was generally stable across a variety of tested parameters.

#### Differential module expression analysis

To test for differences in module gene expression by genotype, we analyzed each module identified in the dataset by plotting the DESeq2 Wald stat, a variance corrected effect size, of its member genes based on pseudo-bulk differential expression analysis. For each module and major cell type (including PCs, which the model was not built with), we compared the distribution of the Wald stat of module genes to the distribution of a randomly sampled gene set of equal size pulled from the grey module. Empirical p-values were calculated from Wilcoxon rank sum test and adjusted across tests using the Benjamini-Hochberg method. Modules with an adjusted p-value < 0.01 were considered significant (DEM).

#### GO enrichment analysis

GO analysis was performed on significant DEGs and DEMs using the *enrichGO* function from the clusterProfiler R package^38^ (version 4.14.6) and org.Mm.eg.db (version 3.20.0), with the gene universe being set to all genes expressed in the cell type the DEGs were identified in or the genes tested in hdWGCNA. Genes and modules were annotated with biological process (BP), cellular component (CC) and molecular function (MF) terms. Terms were deemed significant if they had a corrected p-value < 0.05.

#### Gene set enrichment analysis (GSEA)

Gene set enrichment analysis was performed on the Monocle3 pseudotime gene sets and DEGs from bulk and pseudobulk DE analysis using the gseGO function from the clusterProfiler package with the org.Mm.eg.db database. For the Monocle3 pseudotime gene sets, the significant pseudotime genes were ranked by the difference in expression between the average expression of the gene in cells in the 4^th^ quartile vs 1^st^ quartile of Monocle3 pseudotime scores. For DE contrasts, DEGs were ranked by the DESeq2 Wald stat. Enrichment testing was conducted using the *gseGO* function, significance was assessed using Storey’s q-value (FDR), and terms with FDR < 0.05 were considered significant.

#### Over-representation analysis (ORA)

To determine the significance of overlap between DEGs (P < 0.05) identified in the RNA-seq and external gene sets we used permutation tests. To generate a null distribution, we performed 10,000 permutations. In each iteration, a random set of genes equal in size to the input list was sampled without replacement from the background of genes for DE. The overlap between the custom gene set and each permuted gene set was recorded and a p-value was calculated as the proportion of permutations in which the overlap was greater than or equal to the observed overlap. A Benjamini-Hochberg false discovery rate correction of p-values was applied to account for multiple testing. The ASD gene set was obtained from the SFARI database and Fu et al.^12^ which included only those genes that passed FDR < 0.05 for transmission and de novo association (TADA) for ASD. The epilepsy, microcephaly and macrocephaly gene sets were obtained for the curated DisGeNET database. The CHIP-seq set was obtained from Wade et al.^63^, which included metanalysis of published CHD8 CHIP-seq experiments. We curated our target list to include those that were present across three independent CHIP-seq experiments in Gompers et al.^21^, Platt et al.^22^, and Katayama et al.^23^. The Platt CHIP-seq set was also included as an independent example. The TaDa-seq gene set was derived from ref. Wade et al.^64^.

#### Monocle3 Pseudotime analysis

Developmental trajectories were reconstructed separately for the GC, MLI, and oligodendrocyte lineages using Monocle3^37^. WT and *Chd8^+/5bpdel^* cells were analyzed together, and trajectories were rooted in the corresponding progenitor population. Monocle3 (version 1.3.7) learned a principal graph from the reduced-dimensional representation and assigned each cell a pseudotime value reflecting its position along the inferred developmental trajectory. Genotype differences were evaluated by comparing the median pseudotime of each biological replicate using two-sided Student’s *t*-tests. Genes varying along each trajectory were identified with the graph_test function using the principal graph. This test uses Moran’s I to detect nonrandom spatial patterns of gene expression across the trajectory. Pseudotime-associated genes were classified according to their association with immature or mature states based on their enrichment in the lower vs upper end of pseudotime scores and used in downstream DE and functional enrichment analyses.

### RNA isolation for bulk RNA-sequencing, and bioinformatics analysis

Total RNA was obtained using RNAqueous Total RNA Isolation Kit (cat# AM1912 ThermoFisher Scientifics), and assayed via Agilent RNA 6000 Nano Bioanalyzer kit/instrument. Poly-A-enriched mRNA libraries were prepared at Novogene using Illumina reagents. Libraries were sequenced using Illumina NovaSeq 6000 S4 system, paired-end 150 (PE150) method. Reads were aligned to the mouse genome (GRCm38/mm10) using STAR (version 2.5.4b), and gene counts were produced using featureCounts (version 1.6.0). FeatureCounts processed an average of 37.4 million feature-assigned reads per P12 sample (range, 31.4–47.3 million) and an average of 25.6 million per adult sample (range, 16.2–33.7 million). Data quality was assessed using FastQC^65^ (version 0.11.5), and principal component analysis (PCA) was used to determine presence of sample outliers. Two samples were removed for the analysis based on these criteria. Raw RNA-seq fastq files and a gene count matrix are available on GEO. Genotype and sex were confirmed via expression of sex-linked genes and visualization of the 5bp deletion *Chd8* in IGV. For DE analysis, we used DESeq2 R package^66^ (version 1.46.0). Genes with a minimum of 10 in at least half of the samples were included in the analysis. We included covariates for sex and the first two principal components (PCs) of normalized gene expression, which did not correlate with genotype, were additionally included as covariates in all DESeq2 models. For sex-stratified DE, we excluded the covariate for sex. DESeq2 normalized reads were used for all gene plots.

### *Corin* RNAscope and quantification

To simultaneously detect *Corin* RNA and CALB1 protein in fixed-frozen CB sections, an integrated RNAscope co-detection assay was performed using the RNAscope Multiplex Fluorescent v2 kit (Advanced Cell Diagnostics, ACD) combined with IHC. Briefly, following perfusion, mouse CB were sectioned at 40 μm using a cryostat. Sections were mounted onto charged slides and baked at 60°C for 30 min. Slides were post-fixed in pre-cooled 4% PFA for 60 min at RT, dehydrated through a graded ethanol series (50%, 70%, and 100% EtOH; 5 min each), and air-dried. Endogenous peroxidase activity was quenched with RNAscope Hydrogen Peroxide for 5 min at RT. For antigen retrieval, sections were incubated in boiling 1× Co-Detection Target Retrieval solution (98-102°C) for 5 min, washed with PBS containing 0.1% Tween-20 (PBS-T), and outlined with a hydrophobic barrier pen. Sections were incubated overnight at 4°C in a humidified chamber with rabbit anti-CALB1 antibody (Millipore sigma, 1:200) diluted in Co-Detection Antibody Diluent. On the following day, sections were post-fixed with 4% PFA for 30 min at RT and digested with RNAscope Protease Plus at 40°C for 30 min. *Corin* RNA probes were then hybridized at 40°C for 2 h, followed by sequential signal amplification with AMP 1 (30 min at 40°C), AMP 2 (30 min at 40°C), and AMP 3 (15 min at 40°C), according to the manufacturer’s instructions. Sections were subsequently incubated with RNAscope Multiplex FL v2 HRP-C1 at 40°C for 15 min. TSA Vivid Fluorophore 570 (1:2,000) was applied for 30 min at 40°C, followed by HRP blocking. For CALB1 detection, sections were then incubated with an anti-rabbit Alexa Fluor 488 secondary antibody (1:200) diluted in Co-Detection Antibody Diluent for 30 min at RT. Nuclei were counterstained with DAPI for 3 min, and slides were coverslipped using ProLong Gold Antifade Mountant. Fluorescent signals were visualized and imaged using an Olympus OlyVIA imaging system and an LSM 800 confocal microscope.

ImageJ was used to quantify the *Corin* RNAscope images. First the soma of PCs in all CB lobules were outlined via expression of CALB1. Since RNAscope is a single-molecule FISH protocol we calculated *Corin* expression as the area within the PC soma with detectable fluorescence over background. This value was normalized by the overall area of the PC. A Wilcoxon rank sum test was used to compare the distribution of *Corin* expression within WT and *Chd8^+/5bpdel^*, performed across all cells and within individual regions. A binomial generalized linear mixed-effects model controlling for lobule and sample was used to identify significant differences in the number of PCs expressing *Corin* (any detectable expression) between genotypes.

### Ex vivo electrophysiology

#### *Sli*ce preparation

Mice (P11-12; both sexes) were anesthetized with isoflurane and decapitated. Brains were rapidly removed and immersed in ice-cold sucrose cutting solution (∼315 mOsm; in mM: NaCl 76, NaHCO₃ 25, sucrose 65, D-glucose 25, KCl 2.5, NaH₂PO₄·H₂O 1.4, MgCl₂ 7, CaCl₂ 0.1, sodium ascorbate 0.4, sodium pyruvate 2)^67^. Sagittal CB sections (220 μm thickness) were cut using a vibratome (VT1200S; Leica Microsystems, Wetzlar, Germany) and transferred to an incubation chamber containing oxygenated sucrose cutting solution at 32°C for 30 min. Slices were then transferred to oxygenated artificial cerebrospinal fluid (aCSF ∼315 mOsm; in mM: NaCl 127, NaHCO₃ 25, NaH₂PO₄·H₂O 1.25, KCl 2.5, D-glucose 25, MgCl₂ 1, and CaCl₂ 2) at 32°C for an additional 30 min before being maintained at room temperature (RT) until use. All solutions were continuously bubbled with 95% O₂ and 5% CO₂.

#### Recording

For recordings, slices were mounted onto poly-L-lysine-coated glass coverslips (Sigma-Aldrich, St. Louis, MO, USA) and transferred to a submerged recording chamber superfused with oxygenated aCSF (1-2 mL/min) at RT. Whole-cell patch clamp recordings were obtained from visually identified PCs, selected based on their characteristic location within the PCL and soma morphology, with borosilicate glass pipettes (1.5-2 MΩ; Sutter Instrument Co., Novato, CA, USA). Recordings were performed under a 60× water-immersion objective (1.0 NA) mounted on an Olympus BX51WI microscope. Signals were acquired using a Multiclamp 700B amplifier and a Digidata 1550 digitizer (Molecular Devices, San Jose, CA, USA), digitized at 20 kHz, low-pass filtered at 8 kHz, and recorded using pClamp 11 software (Molecular Devices, San Jose, CA, USA).

#### Intrinsic propertie

Recordings were performed using a potassium-based intracellular solution containing (in mM): K-gluconate 123, KCl 12, HEPES 10, Mg-ATP 4, and Na-GTP 0.3^68,69^. To isolate intrinsic membrane properties, slices were continuously perfused with aCSF containing bicuculline (20 μM, Sigma-Aldrich, St. Louis, MO, USA) and NBQX (10 μM, Abcam, Cambridge, UK) to block GABA_A_ and AMPA receptor-mediated synaptic transmission, respectively. Series resistance and leak current were continuously monitored, and cells were excluded if either parameter changed substantially during the experiment.

Spontaneous activity was recorded for 2 minutes immediately after achieving whole-cell configuration. Basal firing rate was calculated from the first 10 s of recording. Spike train regularity was quantified using interspike interval (ISI)-based metrics, including the coefficient of variation (CV: SD_ISI_/ Mean_ISI_) and CV2 (calculated as: 2 ⋅ ∣ISI_n+1_ − ISI_n_∣ / (ISI_n+1_ + ISI_n_) for each pair of consecutive interspike intervals, and averaged across all intervals.

For passive and active membrane property measurements, membrane potential was maintained at −70 mV. Passive membrane properties were assessed using ten consecutive hyperpolarizing current injections of −10 pA. Input resistance (R_input_) was calculated from the slope of the voltage-current relationship, membrane time constant (τ) from an exponential fit of the voltage response, membrane capacitance (C_m_) as τ/ R_input_, and sag ratio was calculated as the ratio between the steady-state (V_ss_) and peak hyperpolarization responses (V_ss_/V_peak_), where V_ss_ was defined as the mean of the final 30 sampled data points of the hyperpolarizing current step.

Active membrane properties were evaluated using a series of 40 depolarizing current steps delivered in 10 pA increments. Rheobase was defined as the minimum current injection required to evoke an action potential. Action potential threshold was defined as the membrane potential at which *dV/dt* exceeded 20 mV. Action potential amplitude was calculated as the difference between action potential peak and threshold, and half-width was measured at 50% of the action potential amplitude. Rise time and decay time were defined as the time from threshold to peak, and from peak to the fast afterhyperpolarization (fAHP), respectively. Rise slope and decay slope corresponded to the maximum and minimum *dV/dt* values during the upstroke and repolarization phases of the action potential. fAHP was quantified as the difference between baseline membrane potential and the minimum voltage reached within 10 ms following the action potential peak, whereas medium afterhyperpolarization (mAHP) was measured as the difference between baseline membrane potential and the minimum voltage occurring 20-200 ms after the spike. All action potential parameters were quantified from the first action potential elicited at rheobase.

Finally, neuronal gain was estimated from the frequency-current (F-I) relationship using a Hill-type sigmoidal fit. Gain was calculated as the maximum slope of the fitted curve and expressed in Hz/nA^70^. Spike frequency adaptation (SFA) was quantified from the sweep corresponding to 1.5× rheobase using the ratio between the last and first ISI. Definitions and quantification criteria were adapted from previous studies of PC intrinsic electrophysiological properties^71–74^. Analyses were performed using custom scripts in MATLAB*^®^* (MathWorks, Natick, MA, USA). Prior to analysis, line noise contamination was removed using a 60 Hz notch filter.

#### Miniature synaptic currents

Miniature synaptic currents were recorded in voltage-clamp using Cs-based internal solution. For mEPSC recordings, the internal solution contained (in mM): 120 CsMeSO₃, 15 CsCl, 8 NaCl, 10 TEA-Cl, 10 HEPES, 0.5 EGTA, 2 QX-314, 4 Mg-ATP, and 0.3 Na-GTP^75^. For mIPSC recordings, a high-chloride Cs-based internal solution was used containing (in mM): 140 CsCl, 4 NaCl, 0.5 CaCl₂, 5 EGTA, 10 HEPES, 2 Mg-ATP, and 5 QX-314-Cl^76^. Miniature synaptic currents were isolated in the presence of tetrodotoxin (TTX; 1 μM, Abcam, Cambridge, UK) together with either bicuculline (20 μM, Sigma-Aldrich, St. Louis, MO, USA) for mEPSC recordings or NBQX (10 μM, Abcam, Cambridge, UK) for mIPSC recordings. mPSCs were analyzed offline in Igor Pro (v. 7.08; WaveMetrics, Lake Oswego, OR, USA) using custom-written scripts. Event detection was based on threshold crossings in the first derivative of filtered current traces. For each detected event, the local peak and baseline were identified within a ±10 ms window surrounding the detection time, and event amplitude was calculated as the difference between these values. Event times were stored for subsequent analyses of event frequency and inter-event interval (IEI) distributions.

### Cerebellar and PC soma volume analysis

Section areas were quantified using ImageJ (v1.54, National Institutes of Health, Bethesda, MD, USA). For CB volume estimation, one out of every three serial sections spanning the entire cerebellum was analyzed. Total CB volume was calculated using the Cavalieri principle ^77^ according to the following equation:

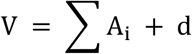

where *V* represents CB volume, *Aᵢ* corresponds to the area of each analyzed section, and *d* is the distance between analyzed sections (240 μm). Volumes are reported in mm³.

Similarly, the somatic volume of biocytin-filled neurons was estimated from confocal image stacks. Soma area was measured across consecutive optical planes, and total somatic volume was calculated by summing the measured areas and multiplying by the distance between adjacent planes.

### PC dendritic morphology analysis

For morphological reconstruction experiments, PCs were filled intracellularly with biocytin (0.05%; 0.5 mg/mL in the internal solution; Life Technologies, Carlsbad, CA, USA) during whole-cell recordings. Following cell filling, the pipette was slowly withdrawn to allow membrane resealing, and slices were maintained for at least 15 min to permit diffusion of biocytin into distal dendritic processes. After recording, slices were fixed overnight in 4% PFA in 0.1 M phosphate buffer. Biocytin-filled neurons were visualized using streptavidin-Alexa Fluor 488 (1:500, Thermo Fisher Scientific, Waltham, MA, USA) in a blocking solution containing 10% normal goat serum (Abcam, Cambridge, UK) and 0.5% Triton X-100 (Sigma-Aldrich, St. Louis, MO, USA) in PBS prior to mounting.

Sholl analysis was performed in ImageJ (v1.54, National Institutes of Health, Bethesda, MD, USA) using the SNT plugin. Images were thresholded and converted to binary masks prior to analysis. The center of the soma was defined as the origin, and concentric circles with a radius increment of 10 μm were generated automatically. The number of dendritic intersections was calculated as the number of times dendrites crossed each concentric circle. Total dendritic length was calculated as the cumulative length of all detected dendritic segments, while the number of dendritic branches corresponded to the total number of branch segments identified within the dendritic arbor.

### Statistical Analysis for electrophysiology and morphology

Statistical analyses were performed using GraphPad Prism 9.3.0 (GraphPad Software, San Diego, CA, USA) or RStudio 2024.12.1 (Posit Software, PBC, Boston, MA, USA). Unless otherwise indicated, comparisons between experimental groups were performed using unpaired two-tailed Student’s *t*-tests or Mann-Whitney. Sholl analysis data (number of intersections as a function of distance from the soma) and frequency-current relationships were analyzed using mixed-effects ANOVA models^78^, with distance or current injection step as the repeated-measures factor and genotype as the between-subject factor. When the assumption of sphericity was violated, degrees of freedom were corrected using the Geisser-Greenhouse correction^79^. Post-hoc pairwise comparisons were corrected for multiple testing using the Benjamini-Hochberg procedure with a false discovery rate (FDR)^80^ of 10%. Cumulative distributions of miniature synaptic event amplitudes and inter-event intervals were compared using the two-sample Kolmogorov-Smirnov test. Data are presented as mean ± SEM and the statistical significance was defined as *p* < 0.05.

## Data availability

The raw sequencing data and processed files generated in this study have been deposited in the NCBI Gene Expression Omnibus (GEO). The bulk RNA-seq data from the P12 and adult cerebellar dissections of constitutive *Chd8^+/5bpdel^*and wildtype littermates, including raw FASTQ files and gene count matrices are available under accession number GSE351236. The single-nucleus RNA-seq data including raw FASTQ files, cell metadata information, and the integrated Seurat object are available under accession number GSE351236. Reads were aligned to the mouse mm10 (GRCm38) reference genome. All other data supporting the findings of this study, including the values underlying the graphs in the main and supplemental figures are provided within the paper and its Supplemental Information, including the supplemental tables.

## Code availability

The code used for the transcriptomic analyses of this study is available at https://github.com/NordNeurogenomicsLab/p12_cb. The code used for the morphological and electrophysiological analyses of this study is available at https://github.com/FioravanteLab/2026_Seban_Chd8-p12.

## Notes

### Competing Interest Statement

The authors have declared no competing interest.

### Summary of Updates

This updated version has been submitted for peer review.

## REFERENCES

1. Rudolph, S. et al. Cognitive-Affective Functions of the Cerebellum. J. Neurosci. 43, 7554– 7564 (2023).

2. Adamaszek, M. et al. Consensus Paper: Cerebellum and Emotion. The Cerebellum 16, 552–576 (2017).

3. Sokolov, A. A., Miall, R. C. & Ivry, R. B. The Cerebellum: Adaptive Prediction for Movement and Cognition. Trends Cogn. Sci. 21, 313–332 (2017).

4. Schmahmann, J. D., Guell, X., Stoodley, C. J. & Halko, M. A. The Theory and Neuroscience of Cerebellar Cognition. Annu. Rev. Neurosci. 42, 337–364 (2019).

5. Limperopoulos, C. et al. Does Cerebellar Injury in Premature Infants Contribute to the High Prevalence of Long-term Cognitive, Learning, and Behavioral Disability in Survivors? Pediatrics 120, 584–593 (2007).

6. Moberget, T. & Ivry, R. B. Prediction, Psychosis, and the Cerebellum. Biol. Psychiatry Cogn. Neurosci. Neuroimaging 4, 820–831 (2019).

7. Reeber, S. L., Otis, T. S. & Sillitoe, R. V. New roles for the cerebellum in health and disease. Front. Syst. Neurosci. 7, (2013).

8. Bloomer, B. F., Morales, J. J., Bolbecker, A. R., Kim, D.-J. & Hetrick, W. P. Cerebellar Structure and Function in Autism Spectrum Disorder. J. Psychiatry Brain Sci. 7, e220003 (2022).

9. Dalal, J. S. et al. Loss of Tsc1 in cerebellar Purkinje cells induces transcriptional and translation changes in FMRP target transcripts. eLife 10, e67399 (2021).

10. Ha, S. et al. Cerebellar Shank2 Regulates Excitatory Synapse Density, Motor Coordination, and Specific Repetitive and Anxiety-Like Behaviors. J. Neurosci. 36, 12129– 12143 (2016).

11. Carey, M. R. The cerebellum. Curr. Biol. 34, R7–R11 (2024).

12. Fu, J. M. et al. Rare coding variation provides insight into the genetic architecture and phenotypic context of autism. Nat. Genet. 54, 1320–1331 (2022).

13. Autism Spectrum Disorder Working Group of the Psychiatric Genomics Consortium et al. Identification of common genetic risk variants for autism spectrum disorder. Nat. Genet. 51, 431–444 (2019).

14. Alotaibi, M. & Ramzan, K. A de novo variant of CHD8 in a patient with autism spectrum disorder. Discoveries 8, e107 (2020).

15. An, Y. et al. De novo variants in the Helicase-C domain of CHD8 are associated with severe phenotypes including autism, language disability and overgrowth. Hum. Genet. 139, 499–512 (2020).

16. Beighley, J. S. et al. Clinical Phenotypes of Carriers of Mutations in CHD8 or Its Conserved Target Genes. Biol. Psychiatry 87, 123–131 (2020).

17. Bernier, R. et al. Disruptive CHD8 Mutations Define a Subtype of Autism Early in Development. Cell 158, 263–276 (2014).

18. Doummar, D. et al. Childhood-onset progressive dystonia associated with pathogenic truncating variants in *CHD8*. Ann. Clin. Transl. Neurol. 8, 1986–1990 (2021).

19. Douzgou, S. et al. The clinical presentation caused by truncating *CHD8* variants. Clin. Genet. 96, 72–84 (2019).

20. Ostrowski, P. J. et al. The *CHD8* overgrowth syndrome: A detailed evaluation of an emerging overgrowth phenotype in 27 patients. Am. J. Med. Genet. C Semin. Med. Genet. 181, 557–564 (2019).

21. Gompers, A. L. et al. Germline Chd8 haploinsufficiency alters brain development in mouse. Nat. Neurosci. 20, 1062–1073 (2017).

22. Platt, R. J. et al. Chd8 Mutation Leads to Autistic-like Behaviors and Impaired Striatal Circuits. Cell Rep. 19, 335–350 (2017).

23. Katayama, Y. et al. CHD8 haploinsufficiency results in autistic-like phenotypes in mice. Nature 537, 675–679 (2016).

24. Canales, C. P. et al. Increased cortical volume without increased neuron number in heterozygous Chd8 mutant mouse cortex. Preprint at 10.1101/2021.01.11.426290 (2021).

25. Chen, X. et al. Deletion of CHD8 in cerebellar granule neuron progenitors leads to severe cerebellar hypoplasia, ataxia, and psychiatric behavior in mice. J. Genet. Genomics 49, 859–869 (2022).

26. Kawamura, A. et al. The autism-associated protein CHD8 is required for cerebellar development and motor function. Cell Rep. 35, 108932 (2021).

27. Chen, V. S. et al. Histology Atlas of the Developing Prenatal and Postnatal Mouse Central Nervous System, with Emphasis on Prenatal Days E7.5 to E18.5. Toxicol. Pathol. 45, 705– 744 (2017).

28. Beekhof, G. C. et al. Differential spatiotemporal development of Purkinje cell populations and cerebellum-dependent sensorimotor behaviors. eLife 10, e63668 (2021).

29. Leto, K. et al. Consensus Paper: Cerebellar Development. The Cerebellum 15, 789–828 (2016).

30. Kozareva, V. et al. A transcriptomic atlas of mouse cerebellar cortex comprehensively defines cell types. Nature 598, 214–219 (2021).

31. Lackey, E. P. et al. Specialized connectivity of molecular layer interneuron subtypes leads to disinhibition and synchronous inhibition of cerebellar Purkinje cells. Neuron 112, 2333–2348.e6 (2024).

32. Hull, C. & Regehr, W. G. The Cerebellar Cortex. Annu. Rev. Neurosci. 45, 151–175 (2022).

33. Consalez, G. G., Goldowitz, D., Casoni, F. & Hawkes, R. Origins, Development, and Compartmentation of the Granule Cells of the Cerebellum. Front. Neural Circuits 14, 611841 (2021).

34. Marques, S. et al. Oligodendrocyte heterogeneity in the mouse juvenile and adult central nervous system. Science 352, 1326–1329 (2016).

35. Vanlandewijck, M. et al. A molecular atlas of cell types and zonation in the brain vasculature. Nature 554, 475–480 (2018).

36. Nicol, P. B. et al. Robust identification of perturbed cell types in single-cell RNA-seq data. Nat. Commun. 15, 7610 (2024).

37. Trapnell, C. et al. The dynamics and regulators of cell fate decisions are revealed by pseudotemporal ordering of single cells. Nat. Biotechnol. 32, 381–386 (2014).

38. Yu, G., Wang, L.-G., Han, Y. & He, Q.-Y. clusterProfiler: an R Package for Comparing Biological Themes Among Gene Clusters. OMICS J. Integr. Biol. 16, 284–287 (2012).

39. Leto, K. et al. Laminar Fate and Phenotype Specification of Cerebellar GABAergic Interneurons. J. Neurosci. 29, 7079–7091 (2009).

40. Sepp, M. et al. Cellular development and evolution of the mammalian cerebellum. Nature 625, 788–796 (2024).

41. Young, K. M. et al. Oligodendrocyte Dynamics in the Healthy Adult CNS: Evidence for Myelin Remodeling. Neuron 77, 873–885 (2013).

42. Bergles, D. E., Roberts, J. D. B., Somogyi, P. & Jahr, C. E. Glutamatergic synapses on oligodendrocyte precursor cells in the hippocampus. Nature 405, 187–191 (2000).

43. Magomedova, L. et al. ARGLU1 is a transcriptional coactivator and splicing regulator important for stress hormone signaling and development. Nucleic Acids Res. 47, 2856– 2870 (2019).

44. Morabito, S., Reese, F., Rahimzadeh, N., Miyoshi, E. & Swarup, V. hdWGCNA identifies co-expression networks in high-dimensional transcriptomics data. *Cell Rep*. Methods 3, 100498 (2023).

45. Satija, R., Farrell, J. A., Gennert, D., Schier, A. F. & Regev, A. Spatial reconstruction of single-cell gene expression data. Nat. Biotechnol. 33, 495–502 (2015).

46. Chen, X. et al. Transcriptomic mapping uncovers Purkinje neuron plasticity driving learning. Nature 605, 722–727 (2022).

48. Yan, W., Wu, F., Morser, J. & Wu, Q. Corin, a transmembrane cardiac serine protease, acts as a pro-atrial natriuretic peptide-converting enzyme. Proc. Natl. Acad. Sci. 97, 8525–8529 (2000).

48. Paulsen, B. et al. Autism genes converge on asynchronous development of shared neuron classes. Nature 602, 268–273 (2022).

49. Yim, K. M. et al. Cell-type-specific dysregulation of gene expression due to Chd8 haploinsufficiency during mouse cortical development. Cell Genomics 5, 100986 (2025).

50. Osório, C. et al. Purkinje cell intrinsic activity shapes cerebellar development and function. Nat. Commun. 17, 3688 (2026).

51. Kelly, E. et al. Regulation of autism-relevant behaviors by cerebellar–prefrontal cortical circuits. Nat. Neurosci. https://doi.org/10.1038/s41593-020-0665-z (2020) doi:10.1038/s41593-020-0665-z.

52. Stoodley, C. J. et al. Altered cerebellar connectivity in autism and cerebellar-mediated rescue of autism-related behaviors in mice. Nat. Neurosci. 20, 1744–1751 (2017).

53. Jahncke, J. N. & Wright, K. M. Tools for *Cre* –Mediated Conditional Deletion of Floxed Alleles from Developing Cerebellar Purkinje Cells. eneuro 11, ENEURO.0149-24.2024 (2024).

54. Suetterlin, P. et al. Altered Neocortical Gene Expression, Brain Overgrowth and Functional Over-Connectivity in Chd8 Haploinsufficient Mice. *Cereb. Cortex* **28**, 2192–2206 (2018).

55. Kano, M. & Hashimoto, K. Activity-Dependent Maturation of Climbing Fiber to Purkinje Cell Synapses during Postnatal Cerebellar Development. The Cerebellum 11, 449–450 (2012).

56. Jung, H. et al. Sexually dimorphic behavior, neuronal activity, and gene expression in Chd8-mutant mice. Nat. Neurosci. 21, 1218–1228 (2018).

57. Hulbert, S. W. et al. A Novel *CHD8* Mutant Mouse Displays Altered Ultrasonic Vocalizations and Enhanced Motor Coordination. Autism Res. 13, 1685–1697 (2020).

58. Niu, X. et al. The Deficiency of the ASD-Related Gene CHD8 Disrupts Behavioral Patterns and Inhibits Hippocampal Neurogenesis in Mice. J. Mol. Neurosci. 74, 103 (2024).

60. Martelotto, L. ‘Frankenstein’ protocol for nuclei isolation from fresh and frozen tissue.

60. Young, M. D. & Behjati, S. SoupX removes ambient RNA contamination from droplet-based single-cell RNA sequencing data. GigaScience 9, giaa151 (2020).

61. McGinnis, C. S., Murrow, L. M. & Gartner, Z. J. DoubletFinder: Doublet Detection in Single-Cell RNA Sequencing Data Using Artificial Nearest Neighbors. Cell Syst. 8, 329–337.e4 (2019).

62. Carter, R. A. et al. A Single-Cell Transcriptional Atlas of the Developing Murine Cerebellum. Curr. Biol. 28, 2910–2920.e2 (2018).

63. Wade, A. A., Lim, K., Catta-Preta, R. & Nord, A. S. Common CHD8 Genomic Targets Contrast With Model-Specific Transcriptional Impacts of CHD8 Haploinsufficiency. Front. Mol. Neurosci. 11, 481 (2019).

64. Wade, A. A. et al. In vivo targeted DamID identifies CHD8 genomic targets in fetal mouse brain. iScience 24, 103234 (2021).

65. Andrews, S. FastQC: a quality control tool for high throughput sequence data. Babraham Bioinformatics (2010).

66. Love, M. I., Huber, W. & Anders, S. Moderated estimation of fold change and dispersion for RNA-seq data with DESeq2. Genome Biol. 15, 550 (2014).

67. McGarry, L. M. & Carter, A. G. Prefrontal Cortex Drives Distinct Projection Neurons in the Basolateral Amygdala. Cell Rep. 21, 1426–1433 (2017).

68. Povysheva, N. V., Zaitsev, A. V., Gonzalez-Burgos, G. & Lewis, D. A. Electrophysiological Heterogeneity of Fast-Spiking Interneurons: Chandelier versus Basket Cells. PLoS ONE 8, e70553 (2013).

69. Cadwell, C. R. et al. Electrophysiological, transcriptomic and morphologic profiling of single neurons using Patch-seq. Nat. Biotechnol. 34, 199–203 (2016).

70. Tang, Y. et al. Modulation of the dynamics of cerebellar Purkinje cells through the interaction of excitatory and inhibitory feedforward pathways. PLOS Comput. Biol. 17, e1008670 (2021).

71. Raman, I. M. & Bean, B. P. Properties of Sodium Currents and Action Potential Firing in Isolated Cerebellar Purkinje Neurons. Ann. N. Y. Acad. Sci. 868, 93–96 (1999).

72. Sacco, T. & Tempia, F. A-Type potassium currents active at subthreshold potentials in mouse cerebellar purkinje cells. J. Physiol. 543, 505–520 (2002).

73. Womack, M. & Khodakhah, K. Active Contribution of Dendrites to the Tonic and Trimodal Patterns of Activity in Cerebellar Purkinje Neurons. J. Neurosci. 22, 10603–10612 (2002).

74. McKay, B. E. & Turner, R. W. Physiological and morphological development of the rat cerebellar Purkinje cell. J. Physiol. 567, 829–850 (2005).

75. Bateup, H. S., Takasaki, K. T., Saulnier, J. L., Denefrio, C. L. & Sabatini, B. L. Loss of Tsc1 In Vivo Impairs Hippocampal mGluR-LTD and Increases Excitatory Synaptic Function. J. Neurosci. 31, 8862–8869 (2011).

76. Houston, C. M., Bright, D. P., Sivilotti, L. G., Beato, M. & Smart, T. G. Intracellular Chloride Ions Regulate the Time Course of GABA-Mediated Inhibitory Synaptic Transmission. J. Neurosci. 29, 10416–10423 (2009).

77. Hosseini-Sharifabad, M. & Sabahi, A. Stereological estimation of granule cell number and purkinje cell volume in the cerebellum of noise-exposed young rat. Iran. J. Med. Sci. 39, 387–390 (2014).

78. Scheffé, H. The Analysis of Variance. (Wiley-Interscience, New York, 1999).

79. Geisser, S. & Greenhouse, S. W. An Extension of Box’s Results on the Use of the $F$ Distribution in Multivariate Analysis. Ann. Math. Stat. 29, 885–891 (1958).

80. Benjamini, Y. & Hochberg, Y. Controlling the False Discovery Rate: A Practical and Powerful Approach to Multiple Testing. J. R. Stat. Soc. Ser. B Stat. Methodol. 57, 289–300 (1995).

