## Supplementary figures for "Precocious maturation and Purkinje cell dysfunction in the mouse *Chd8* haploinsufficient cerebellum"

### SUPPLEMENTAL FIGURES

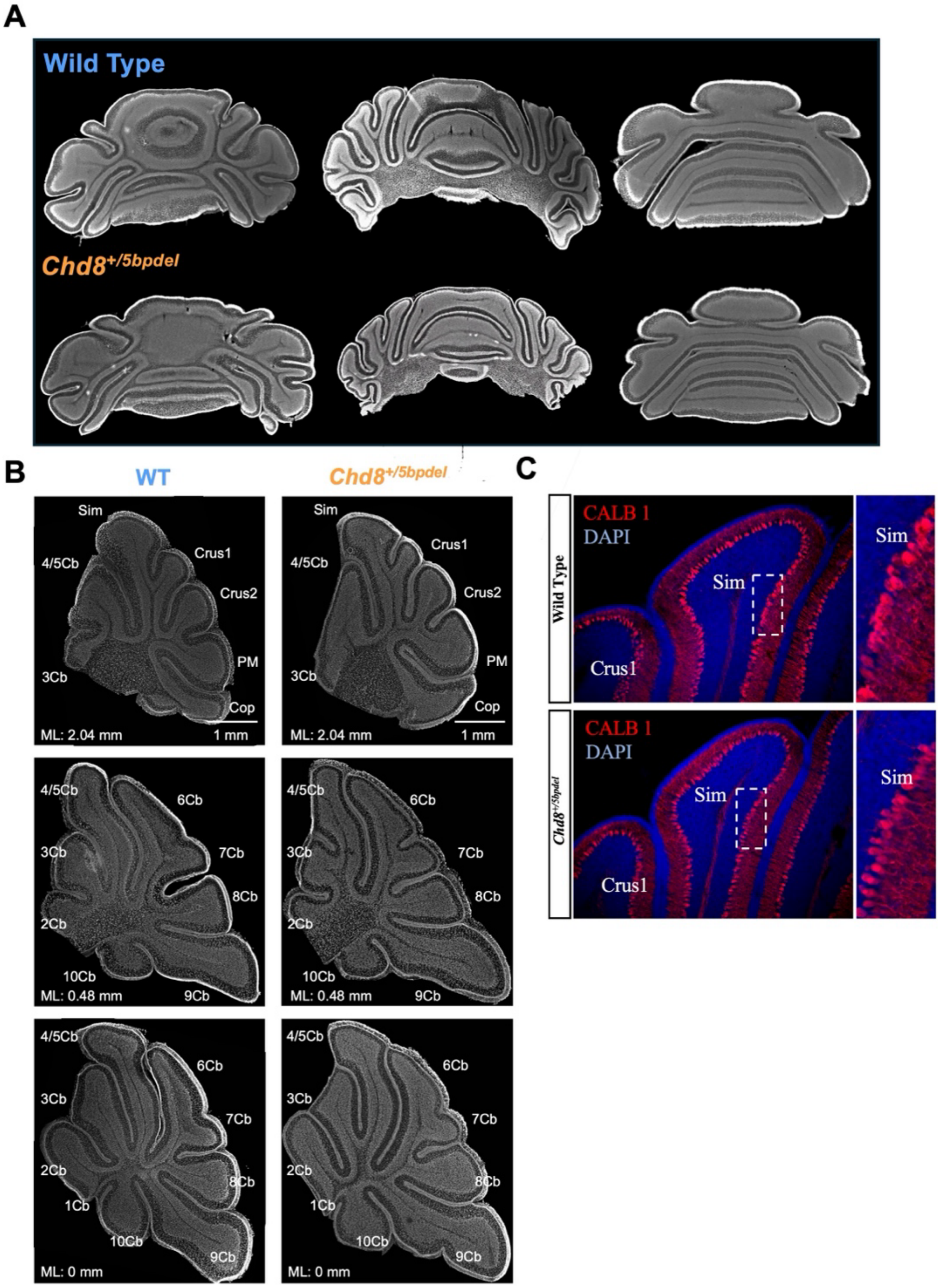


**Supplemental Figure 1. *Chd8* haploinsufficient mice display no gross cerebellar malformations or disruption of Purkinje cell monolayer organization. A**) Representative DAPI-stained serial coronal cerebellar sections from wild-type (WT) and Chd8$^{+/5bpdel}$ (HET) mice used to assess gross cerebellar anatomy. **B**) Representative DAPI-stained sagittal sections from the cerebellar hemisphere, paravermis, and vermis of WT and HET mice used to assess cerebellar morphology and volume. ML indicates the mediolateral position of each section. Scale bars, 1 mm. **C**) Representative sagittal sections from WT and HET mice stained with DAPI and an antibody against CALB1 to label Purkinje cells (PCs). Within a region of interest in the simplex (Sim) lobule, PCs were classified as exhibiting monolayer or multilayer organization. The frequency of multilayer organization was not significantly associated with genotype with a Fisher's exact test (P = 0.443). Dashed boxes indicate the regions shown at higher magnification.

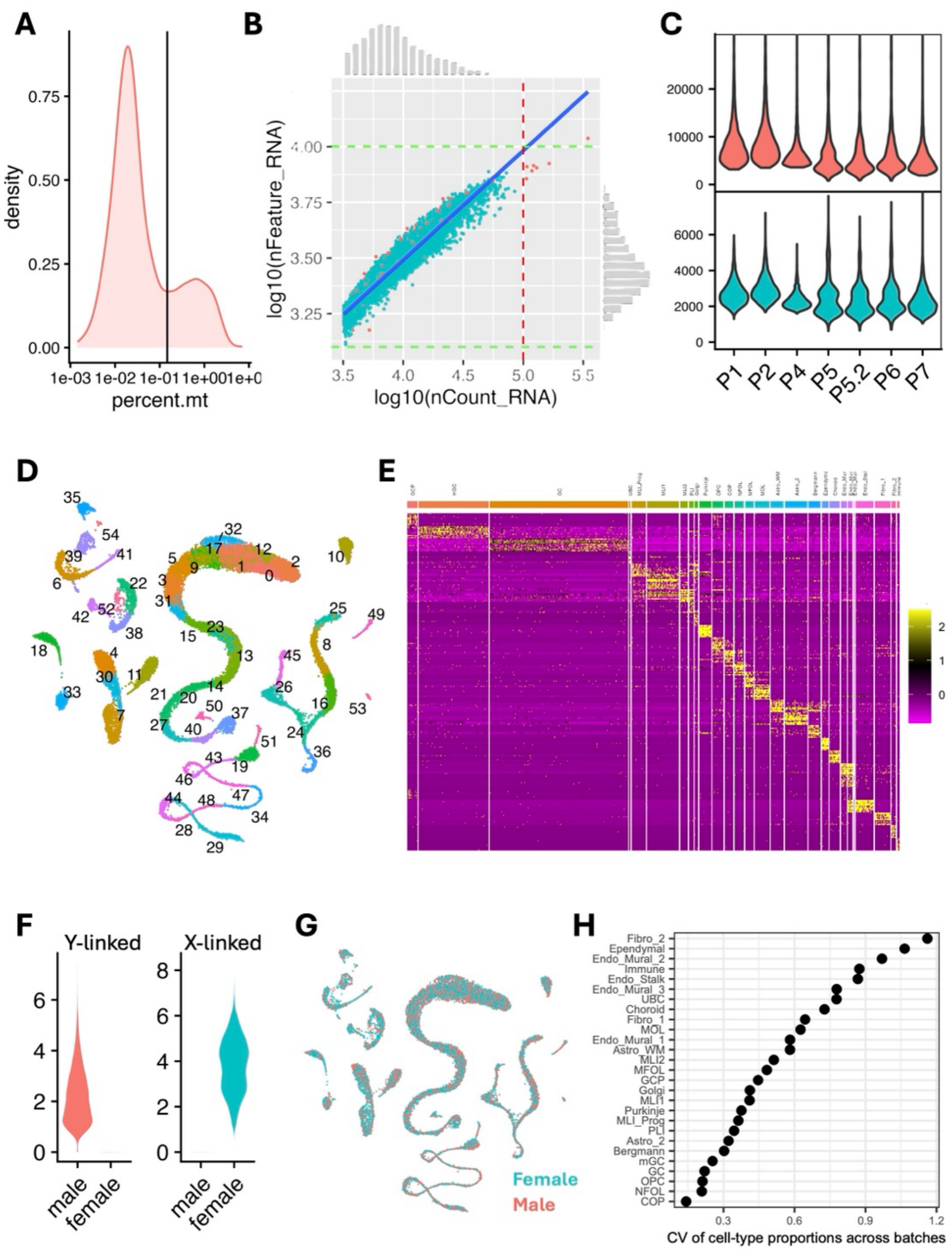


**Supplemental Figure 2. Quality control and annotation of the snRNA-seq dataset.** **A**) Density plot illustrating data-driven identification of the antinode in the percentage of mitochondrial RNA for Pool 1. The antinode was used to establish a pool-specific threshold for excluding low-quality nuclei. **B**) $Log_{10}$-transformed scatterplot of the count-based quality-control metrics, nCount_RNA and nFeature_RNA. Dashed lines indicate filtering thresholds; retained nuclei are shown in cyan, whereas nuclei excluded by count- or mitochondrial-based criteria are shown in red. Marginal histograms show the distribution of each metric. **C**) Violin plots showing nCount_RNA (top) and nFeature_RNA (bottom) across snRNA-seq pools after quality-control filtering. **D**) UMAP representation of data-driven clustering, with nuclei colored and labeled by initial cluster assignment. Clusters assigned to the same cell type were subsequently combined for downstream analyses. **E**) Heatmap showing the top marker genes identified across cell types using Seurat's FindAllMarkers function. **F**) Violin plots showing the expression of Y- and X-linked genes in male and female nuclei from Pool 1. Sex-linked gene expression was used to assign nuclei to male or female samples and facilitate genotype demultiplexing within each pool. **G**) UMAP colored by inferred sex, demonstrating representation of both sexes across clusters. **H**) Coefficient of variation (CV) in the proportion of each cell type across snRNA-seq pools.

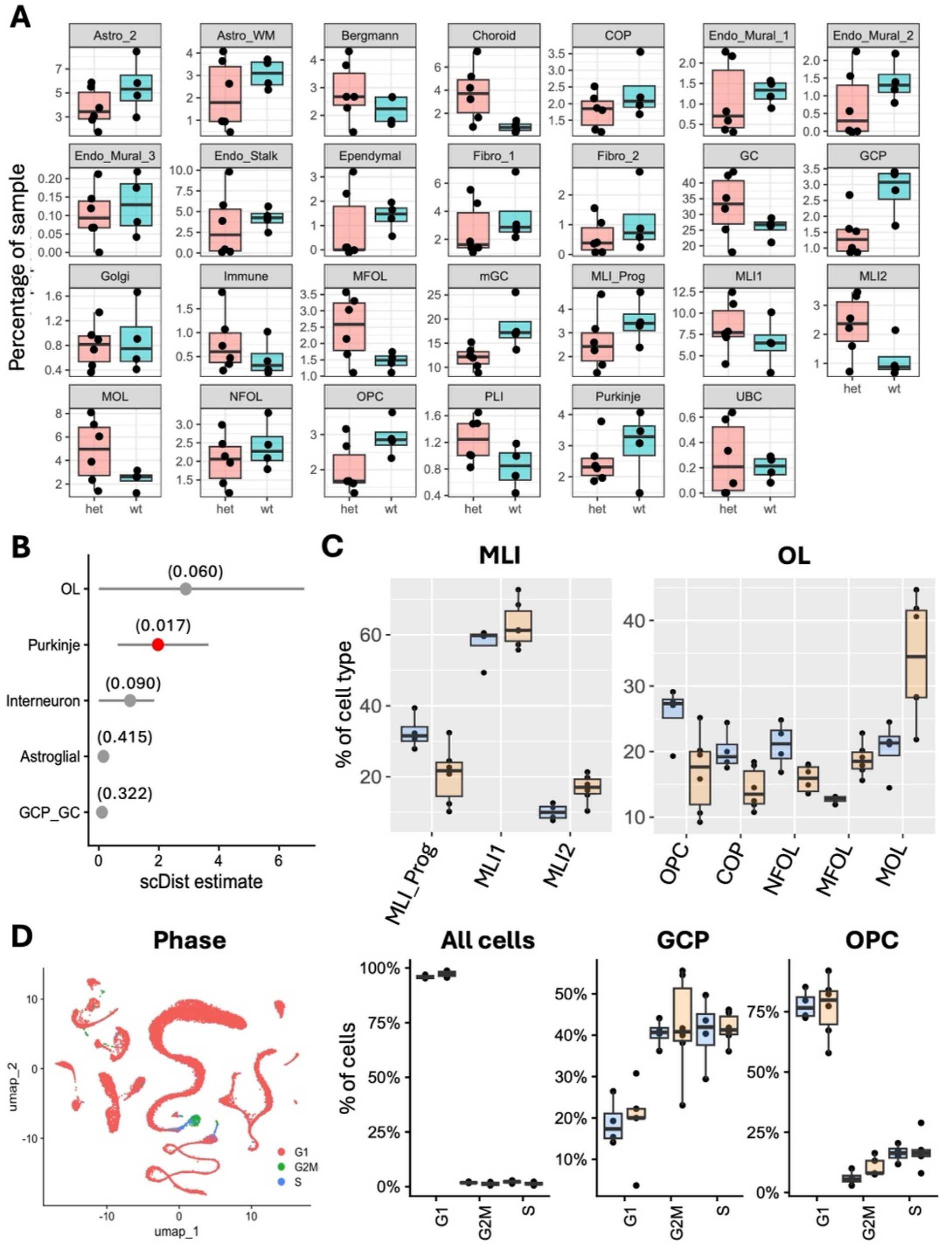


**Supplemental Figure 3. Differences in cell-type composition and transcriptional state in the snRNA-seq dataset. A**) Relative abundance of each cerebellar cell type in wild-type (WT) and Chd8$^{+/5bpdel}$ (HET) samples. Each point represents one biological sample. Genotype differences were assessed using unpaired Student's *t*-tests. **B**) scDist estimates of the magnitude of genotype-associated transcriptional differences within each major cell type. Points represent the estimated distance between WT and HET transcriptional states, and error bars denote 95% posterior credible intervals. Nominal P values are shown in parentheses; the red point indicates the cell type that remained significant after false-discovery-rate correction. **C**) Relative proportions of developmental states within the molecular layer interneuron (MLI) and oligodendrocyte (OL) lineages in WT and HET samples. Each state is shown as a percentage of its respective lineage. Genotype differences were assessed using unpaired Student's *t*-tests. **D**) UMAP showing the cell-cycle phase assigned to each nucleus based on the expression of cell-cycle-associated genes. Boxplots show the percentages of nuclei assigned to G1, G2/M, and S phases across all cells and within the proliferative granule cell progenitor (GCP) and oligodendrocyte precursor cell (OPC) populations. Genotype differences were assessed using unpaired Student's *t*-tests.

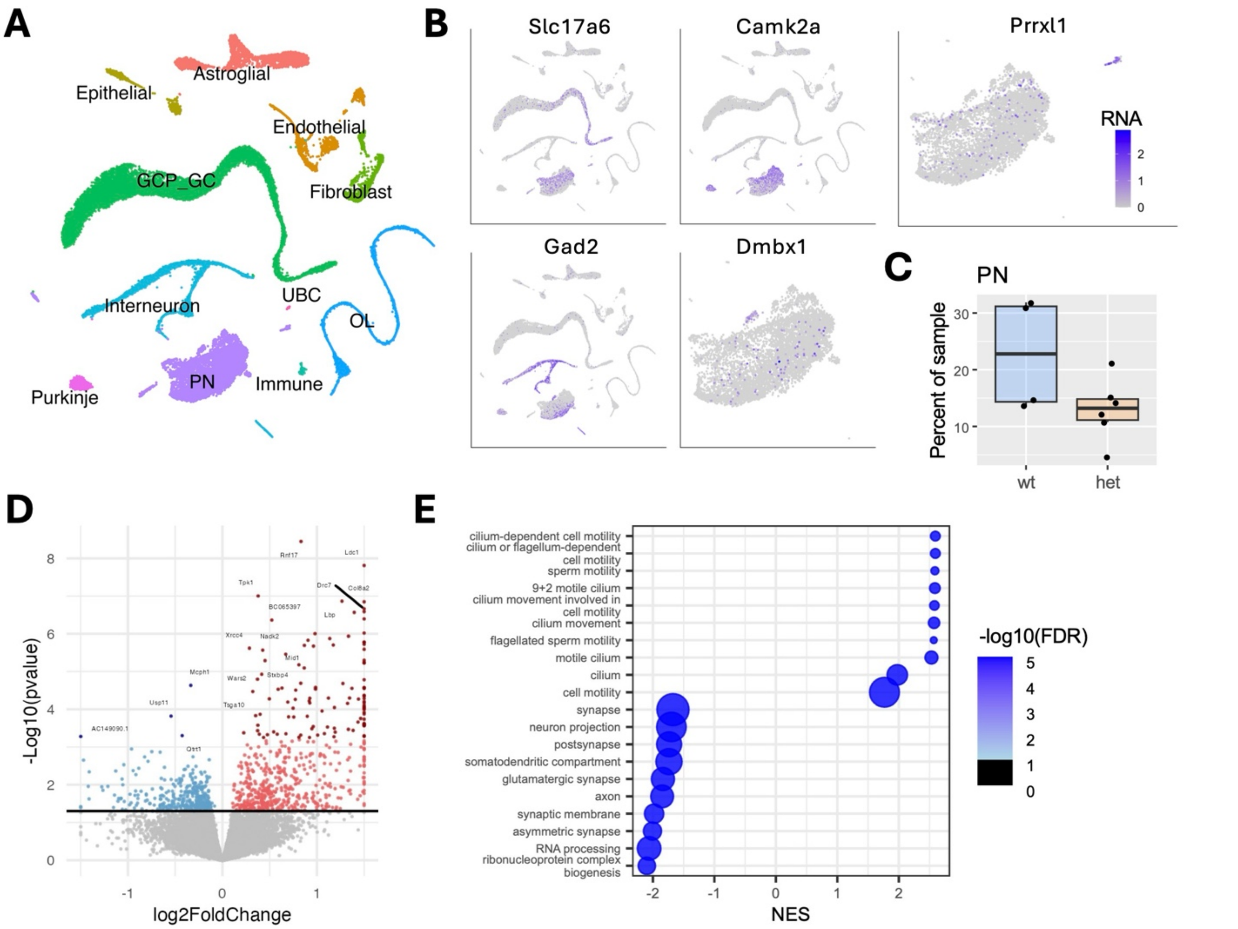


**Supplemental Figure 4. Effects of *Chd8* haploinsufficiency in brainstem neurons.** **A**) UMAP showing cerebellar cortical cell types and brainstem neurons excluded from the primary cerebellar analysis. Three small satellite clusters were tentatively assigned to the dorsal cochlear, trigeminal, and reticular nuclei. The remaining brainstem neurons comprised a broad population of glutamatergic and GABAergic projection neurons (PNs). **B**) Feature plots showing expression of genes used to characterize the PN population: *Slc17a6* (glutamatergic neurons), *Gad2* (GABAergic neurons), *Camk2a* (broad projection-neuron identity), *Dmbx1* (inhibitory deep cerebellar nuclear neurons), and *Prrxl1* (trigeminal neurons). Color intensity indicates normalized RNA expression. **C**) Percentages of PNs in wild-type (WT) and $Chd8^{+/5bpdel}$ (HET) samples. Each point represents one biological sample, and the genotype difference was assessed using an unpaired Student's *t*-test. **D**) Volcano plot showing pseudobulk differential-expression results for PNs, calculated using DESeq2. Light red and light blue indicate nominally significant upregulated and downregulated genes, respectively ($P < 0.05$); dark red and dark blue indicate upregulated and downregulated genes passing an FDR threshold of 0.1. **E**) Representative Gene Ontology (GO) terms identified by gene-set enrichment analysis (GSEA) of PN differential expression. The x-axis indicates the normalized enrichment score (NES), with positive and negative values representing enrichment among upregulated and downregulated genes, respectively. Dot size represents gene-set size, and color indicates enrichment significance.

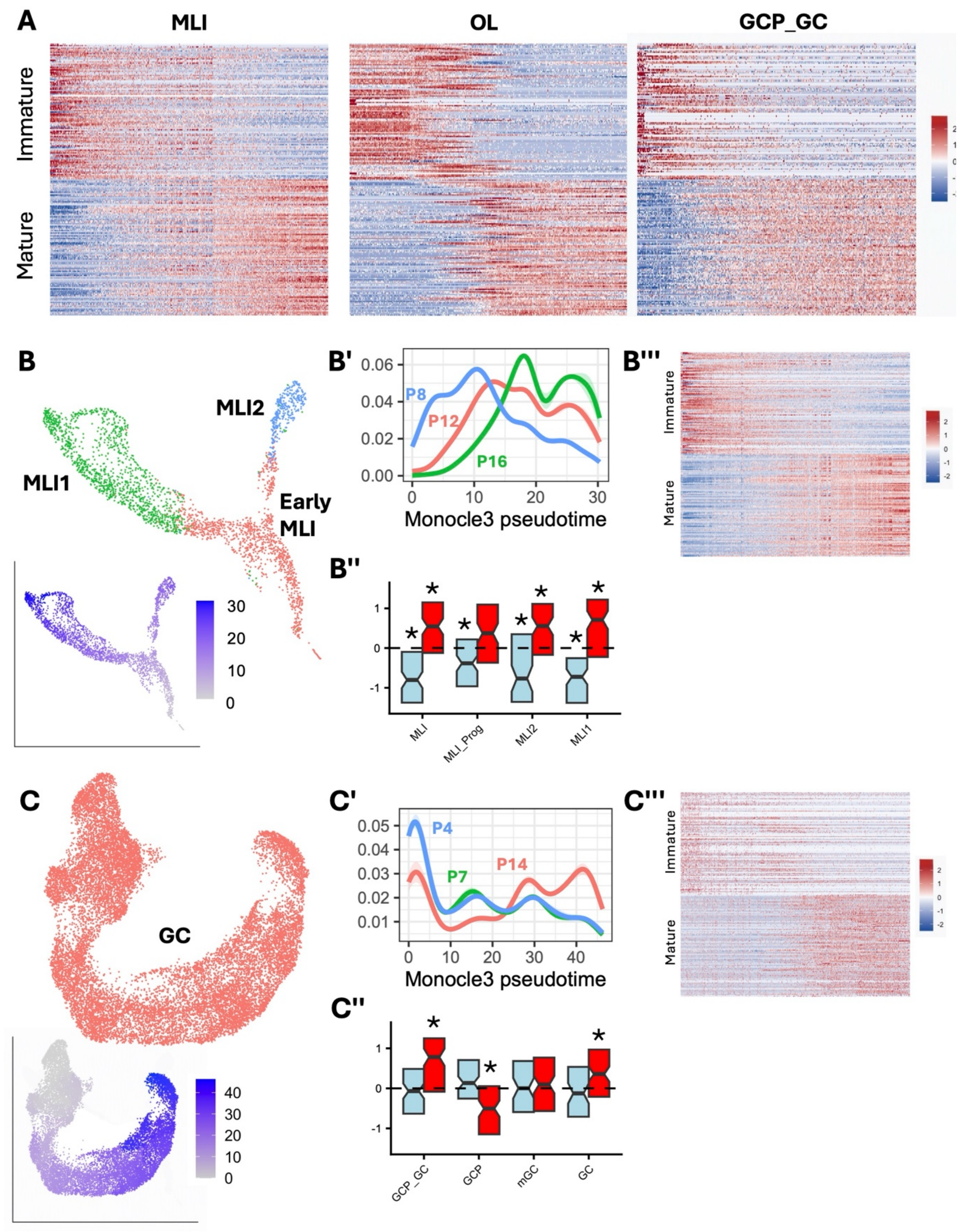


**Supplemental Figure 5. Validation of Monocle3 pseudotime-associated transcriptional signatures in independent snRNA-seq datasets. A**) Heatmaps showing row-scaled expression of the top Monocle3-derived pseudotime-associated genes in the molecular layer interneuron (MLI), oligodendrocyte (OL), and granule cell progenitor-–granule cell (GCP_GC) lineages. Genes are grouped into immature and mature signatures, and nuclei are ordered by increasing pseudotime. **B–B‴**) Validation of the MLI pseudotime signature using longitudinal P8, P12, and P16 snRNA-seq data from Kozareva et al. **B**) UMAP labeled by MLI developmental state, with the inset showing Monocle3 pseudotime values. **B′**) Density distributions of pseudotime values at each developmental age. **B″**) Distributions of DESeq2 Wald statistics in our dataset for genes associated with low (immature; blue) or high (mature; red) pseudotime in the Kozareva et al. MLI trajectory. Asterisks indicate significant differences between gene sets (Wilcoxon rank-sum test, FDR < 0.05). **B‴**) Heatmap showing row-scaled expression of the top pseudotime-associated genes in the Kozareva et al. MLI dataset, with nuclei ordered by increasing pseudotime. **C–C‴**) Validation of the granule cell pseudotime signature using longitudinal P4, P7, and P14 snRNA-seq data from Sarropoulos et al. **C**) UMAP labeled by granule cell identity, with the inset showing Monocle3 pseudotime values. **C′**) Density distributions of pseudotime values at each developmental age. **C″)** Distributions of DESeq2 Wald statistics in our dataset for genes associated with low (immature; blue) or high (mature; red) pseudotime in the Sarropoulos et al. granule cell trajectory. Asterisks indicate significant differences between gene sets (Wilcoxon rank-sum test, FDR < 0.01). **C‴**) Heatmap showing row-scaled expression of the top pseudotime-associated genes in the Sarropoulos et al. granule cell dataset, with nuclei ordered by increasing pseudotime.

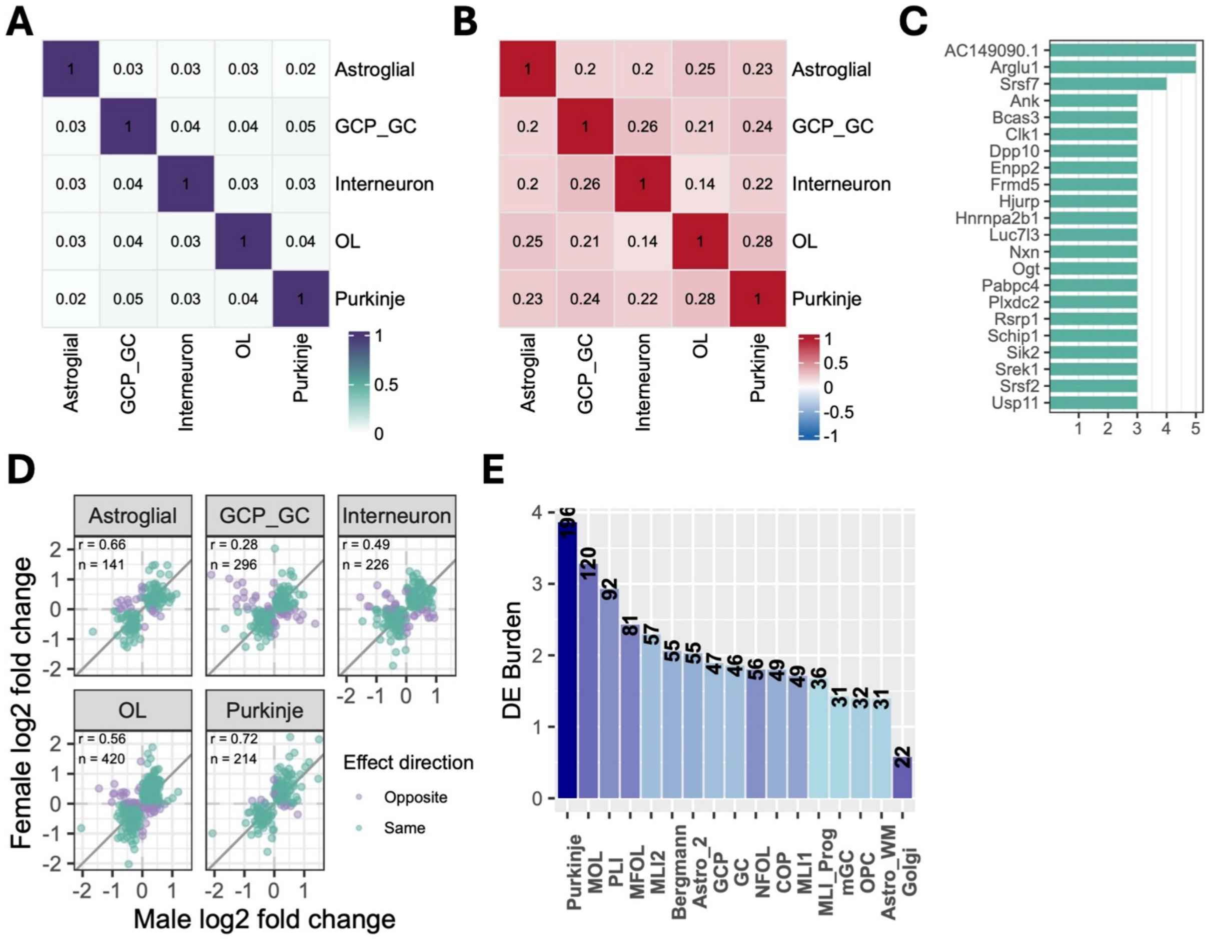


**Supplemental Figure 6. Cross-cell-type and sex-specific comparisons of differential expression in the snRNA-seq dataset. A**) Similarity matrix showing the overlap of differentially expressed genes (DEGs) ($P < 0.05$) between major cerebellar cell types. **B**) Pairwise Pearson correlation coefficients for DEG effect sizes across cell types. **C**) DEGs shared most frequently across cell types. Bar length indicates the number of cell types in which each gene was differentially expressed. **D**) Concordance of genotype-associated $\log_2$ fold changes between male and female samples within each cell type. Each point represents a gene and is colored according to whether its effect direction was concordant or discordant between sexes. Pearson's correlation coefficient (*r*) and the number of genes compared (*n*) are shown in each panel. **E**) Differential-expression burden after downsampling to a maximum of 20 cells per biological sample within each cell type. DE burden was calculated as the percentage of expressed genes with $P < 0.05$; values above the bars indicate the number of DEGs identified.

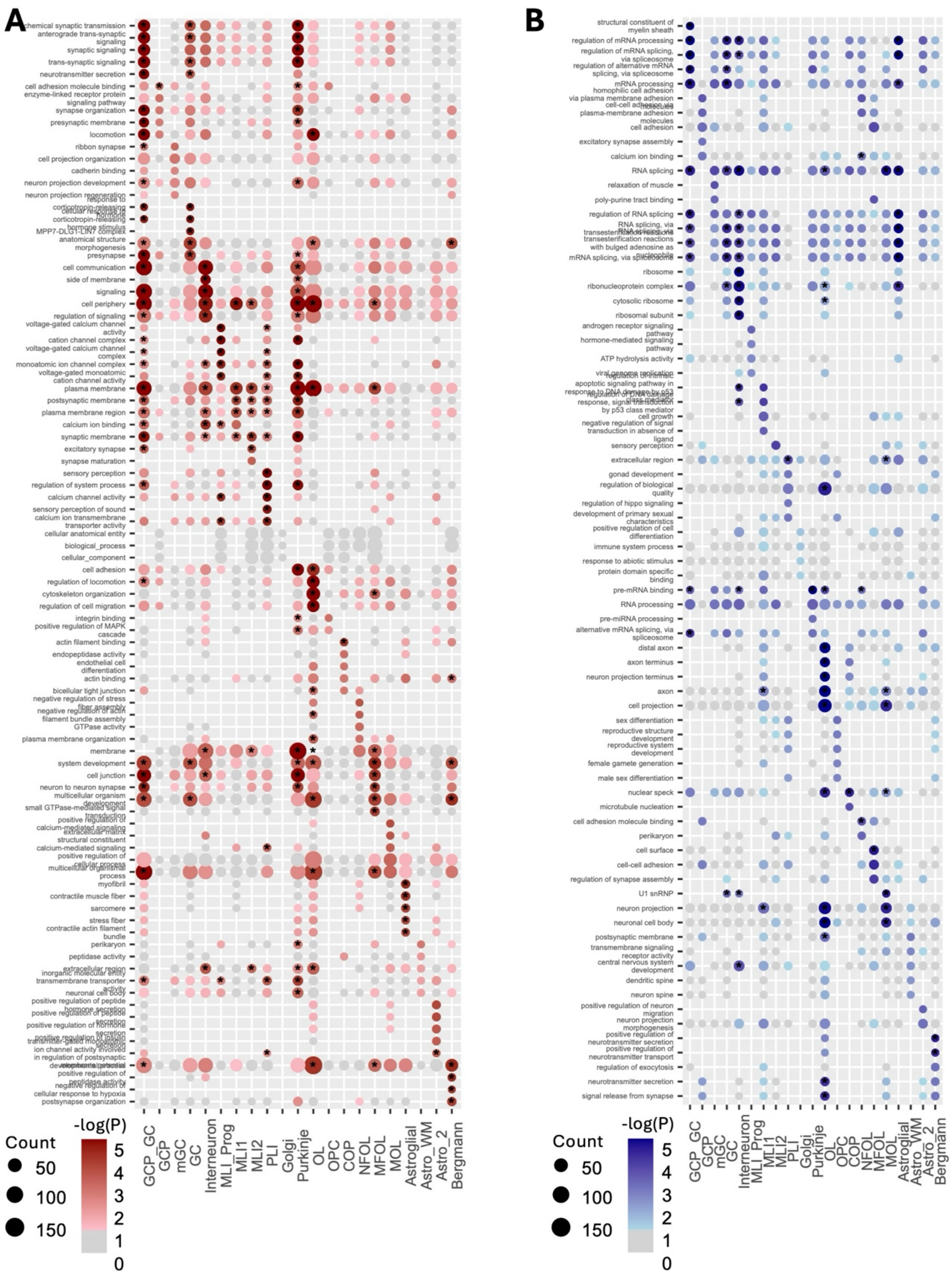


**Supplemental Figure 7: Cell type-specific functional enrichment of differentially expressed genes.** Gene Ontology (GO) overrepresentation analysis of **A**) upregulated and **B**) downregulated genes identified within each cerebellar cell type (P < 0.05). Dot size represents the number of DEGs associated with each GO term, and color intensity represents the significance of enrichment. Gray dots indicate terms without significant enrichment (P > 0.1), and asterisks indicate terms passing the specified multiple-testing threshold (FDR < 0.1). Top 5 (by significance) enriched GO terms per sample are shown.

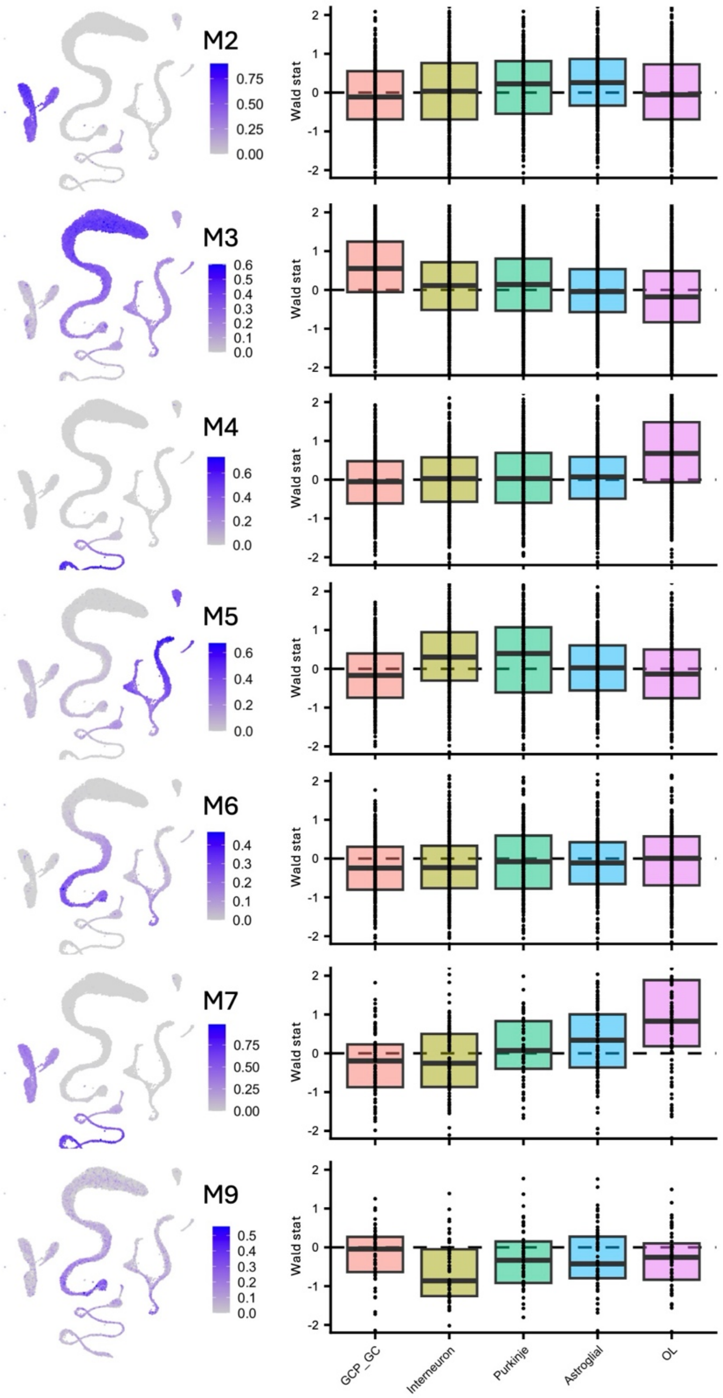


**Supplemental Figure 8. Expression and differential regulation of additional hdWGCNA modules across cerebellar cell types.** Feature plots show the relative expression of hdWGCNA modules M2–M7 and M9 across the snRNA-seq UMAP, as calculated using Seurat's AddModuleScore function. Corresponding boxplots show the distributions of DESeq2 Wald statistics for genes belonging to each module across the major cerebellar cell types. Positive and negative Wald statistics indicate higher and lower expression, respectively, in *Chd8*$^{+/5bpdel}$ mice relative to wild-type controls. Dashed horizontal lines indicate the median Wald statistic of genes in the unassigned gray module for the corresponding cell type. Boxes represent the interquartile range with the median shown by the center line; whiskers extend to 1.5 times the interquartile range.

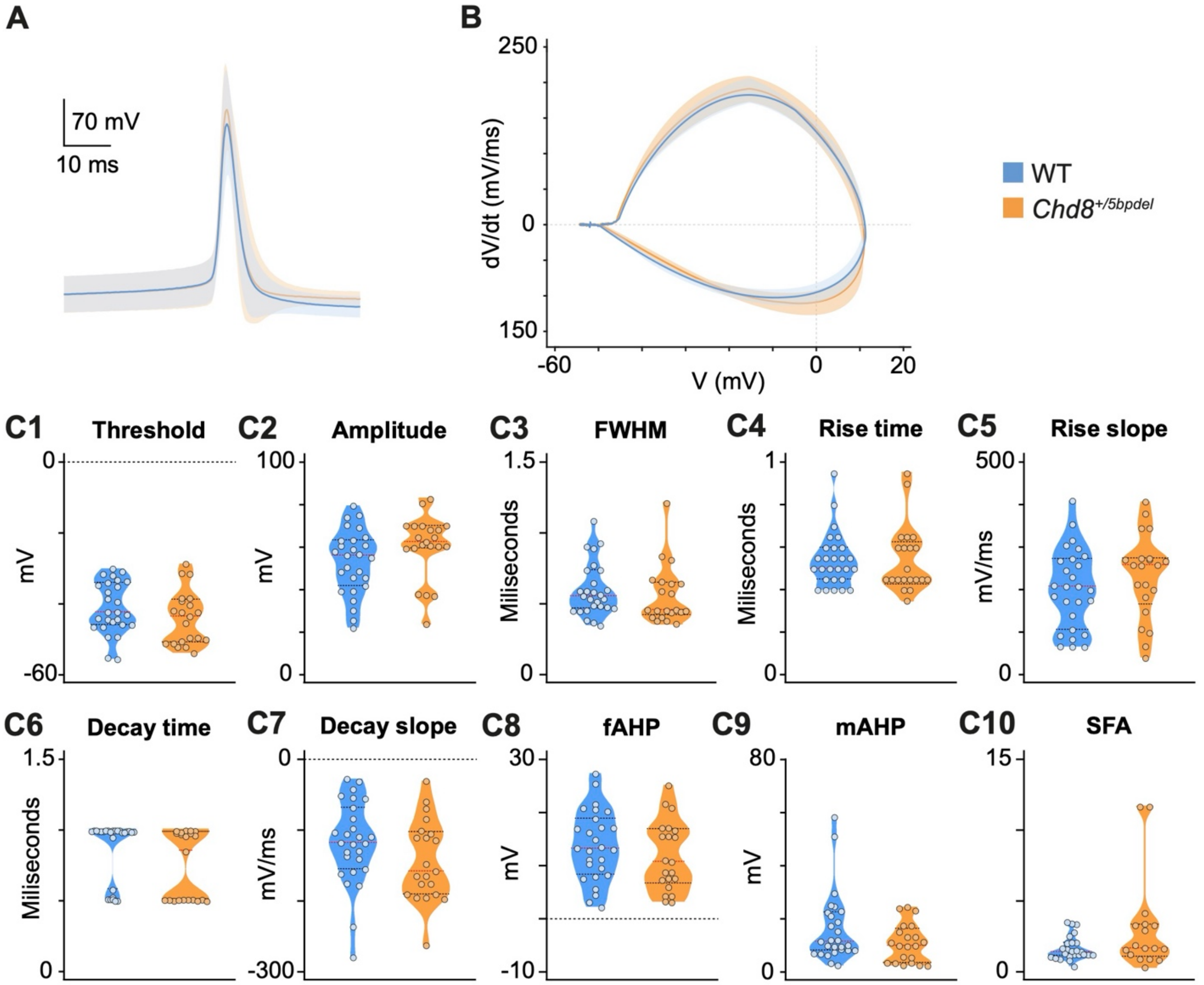


**Supplemental Figure 9: Active membrane properties of Purkinje cells are unaltered in *Chd8$^{+/5bpdel}$* mice. A**) Average waveform of the first action potential (AP) elicited at rheobase in Purkinje cells from WT (blue) and *Chd8$^{+/5bpdel}$* (orange) mice. **B**) Phase plots (dV/dt versus membrane potential) of the APs shown in A. Mean ± SEM is shown. **C1-C10**) Quantification of AP properties: **C1**) threshold, **C2**) amplitude, **C3**) full width at half maximum (FWHM), **C4**) rise time, **C5**) maximum rise slope, **C6**) decay time, **C7**) maximum decay slope, **C8**) fast afterhyperpolarization (fAHP), **C9**) medium afterhyperpolarization (mAHP), and **C10**) spike frequency adaptation (SFA). Red lines in violin plots: median; black dashed lines: 1st and 3rd quartiles. No significant differences were detected between genotypes for any parameter (unpaired Student's t-test or Mann Whitney, $p > 0.05$).

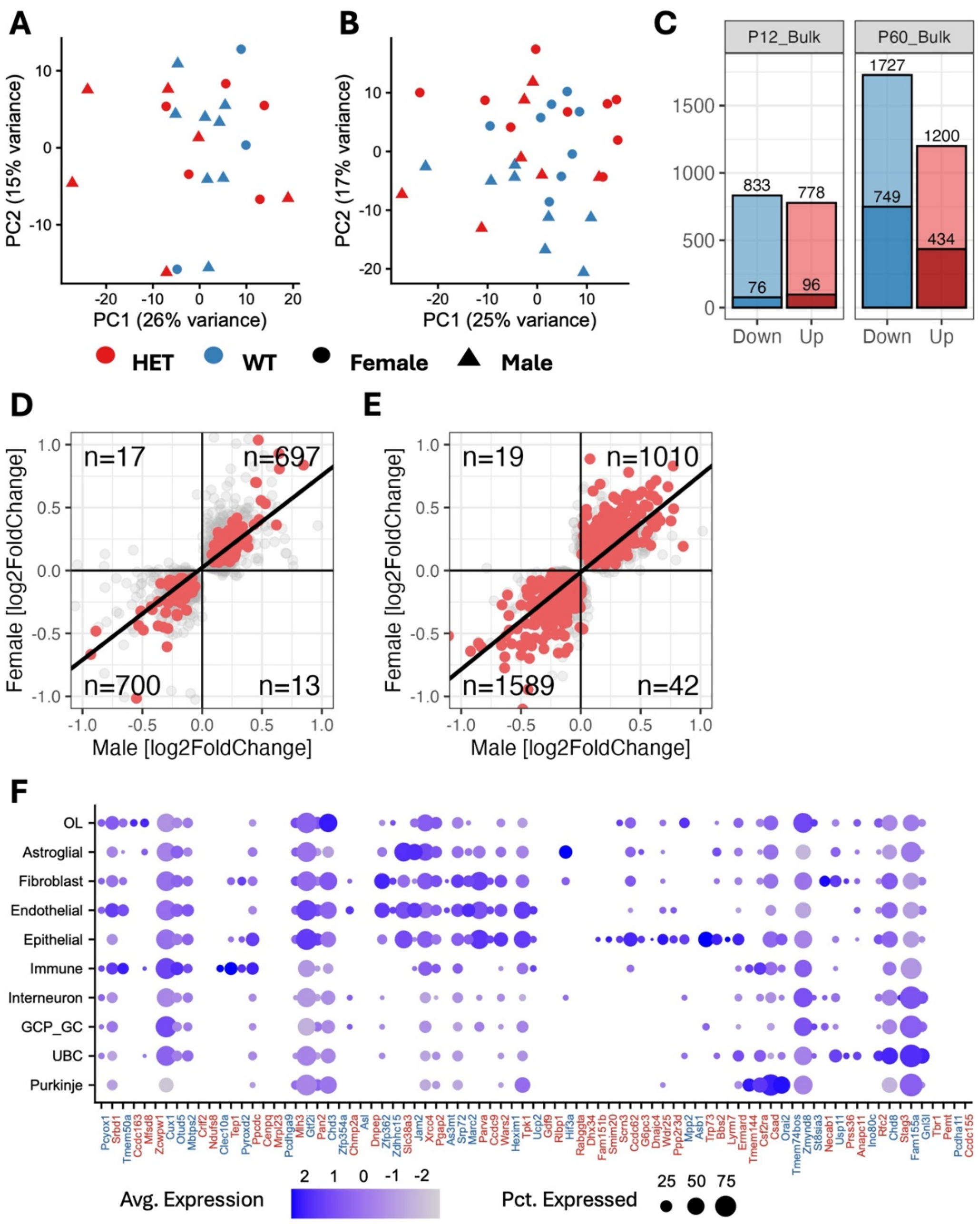


**Supplemental Figure 10: Bulk RNA-seq.** Principal-component analysis (PCA) of cerebellar bulk RNA-sequencing samples collected at **A**) P12 and **B**) P60. Points are colored by genotype and shaped by sex. Axis labels indicate the percentage of variance explained by each principal component. **C**) Numbers of downregulated and upregulated genes at P12 and P60. Full bars represent genes with nominal P < 0.05, whereas the darker portions represent genes passing an FDR threshold of 0.1. **D-E**) Concordance of genotype-associated effect size between male and female samples at **D**) P12 and **E**) P60. Each point represents a gene; gray points denote genes with P < 0.05 and red points denote genes with FDR < 0.1. Quadrant labels indicate the number of genes with concordant or discordant effect directions between sexes, and the diagonal line denotes equal effect sizes in males and females. **F**) Expression of P12 bulk RNA-sequencing DEGs (FDR < 0.1) across cell types in the P12 snRNA-sequencing dataset. Genes expressed in fewer than 10% of nuclei were excluded. Dot size represents the percentage of nuclei expressing each gene, dot color represents scaled average expression, and gene-label color indicates the direction of differential expression in the bulk dataset.

### SUPPLEMENTAL TABLE DESCRIPTIONS

**Supplemental Table 1. Transcriptomics Statistics.** Information on sample size, QC, and specific methods used in Figures 1-4 and 8.

**Supplemental Table 2. Pseudotime gene sets.** Pseudotime based DE assignment for granule cell, molecular layer interneuron, and oligodendrocyte lineages

**Supplemental Table 3. snRNA-seq DE.** Differential expression results (pseudobulk with DESeq2) from single nuclei RNA-seq for all P12 cerebellar cell types and all genes.

**Supplemental Table 4. snRNA-seq GSEA.** GSEA rank-based differential expression functional enrichment (GO terms) for all P12 cerebellar cell types

**Supplemental Table 5. snRNA-seq GO.** DEG functional enrichment (GO terms) for all P12 cerebellar cell types.

**Supplemental Table 6. hdWGCNA modules.** snRNA-seq gene to module assignment from hdWGCNA.

**Supplemental Table 7. PC Electrophysiology Statistics.** Information on sample sizes and statistics for Figures 5-7 and Supplemental Figure 9**.**

**Supplemental Table 8. Bulk RNA-seq DE.** Differential expression results (DESeq2) from bulk RNA-seq for P12 and Adult cerebellum.

**Supplemental Table 9. Bulk GSEA.** GSEA rank-based differential expression functional enrichment (GO terms) for all bulk P12 and adult cerebellum.
